# Comprehensive Evaluation of Diverse Massively Parallel Reporter Assays to Functionally Characterize Human Enhancers Genome-wide

**DOI:** 10.1101/2025.03.25.645321

**Authors:** Junke Zhang, Alden King-Yung Leung, Yutong Zhu, Li Yao, Avery Willis, Xiuqi Pan, Abdullah Ozer, Zhou Zhou, Keith Siklenka, Alejandro Barrera, Jin Liang, Nathaniel D. Tippens, Timothy E. Reddy, John T. Lis, Haiyuan Yu

## Abstract

Massively parallel reporter assays (MPRAs) and self-transcribing active regulatory region sequencing (STARR-seq) have revolutionized enhancer characterization by enabling high-throughput functional assessment of regulatory sequences. Here, we systematically evaluated six MPRA and STARR-seq datasets generated in the human K562 cell line and found substantial inconsistencies in enhancer calls from different labs that are primarily due to technical variations in data processing and experimental workflows. To address these variations, we implemented a uniform enhancer call pipeline, which significantly improved cross-assay agreement. While increasing sequence overlap thresholds enhanced concordance in STARR-seq assays, cross-assay consistency in LentiMPRA was strongly influenced by assay-specific factors. Notably, our results show that LentiMPRA exhibits a strong preference for promoter-associated sequences rather than enhancers. Functional validation using candidate cis-regulatory elements (cCREs) confirmed that epigenomic features such as chromatin accessibility and histone modifications are strong predictors of enhancer activity. Importantly, our study validated transcription as a critical hallmark of active enhancers, demonstrating that highly transcribed regions exhibit significantly higher active rates across assays. Furthermore, we show that transcription enhances the predictive power of epigenomic features, enabling more accurate and refined enhancer annotation. Our study provides a comprehensive framework for integrating different enhancer datasets and underscores the importance of accounting for assay-specific biases when interpreting enhancer activity. These findings refine enhancer identification using massively parallel reporter assays and improve the functional annotation of the human genome.

## Introduction

Enhancers are key cis-regulatory DNA elements that drive transcriptional activity and play a pivotal role in gene regulation. Their influence extends beyond individual gene expression, shaping broader regulatory networks that control cell identity and function. Variants within enhancers have been strongly implicated in complex traits and diseases, emphasizing the importance of systematically identifying and characterizing enhancers to elucidate their contributions to gene expression and disease mechanisms^1–3^.

Traditional reporter gene assays have long been used to characterize enhancer activity by positioning candidate sequences upstream or downstream of a minimal promoter linked to a reporter gene^4–6^. However, enhancers present a much greater challenge for functional characterization than the ∼25,000 protein-coding genes in the human genome due to their vast numbers, sequence variability, and highly context-dependent activity^7,8^ While these traditional reporter gene assays remain functional, the advent of high-throughput sequencing technologies has revolutionized enhancer studies, enabling massively parallel reporter assays (MPRAs) and self-transcribing active regulatory region sequencing (STARR-seq) to profile the regulatory activity of millions of sequences simultaneously^9–11^. These innovations have dramatically expanded our ability to interrogate enhancers on a genome-wide scale, addressing the limitations of conventional low-throughput approaches.

MPRAs, which utilize synthesized oligonucleotide libraries, position candidate sequences upstream of a minimal promoter and tag them with unique barcodes in the 3′ or 5′ UTR of the reporter gene. Regulatory activity is inferred by sequencing RNA transcripts associated with these barcodes^9,10^. Despite their robustness, MPRAs face challenges in testing long DNA sequences and complex libraries due to synthesis and cost limitations^6,12^. Additionally, placing candidate sequences upstream of a promoter may inadvertently capture promoter rather than enhancer activity, confounding the interpretation of regulatory function^13^.

STARR-seq overcomes some of these constraints by placing candidate sequences within the 3′ UTR of a reporter gene, allowing them to self-transcribe and directly quantify enhancer activity based on transcript abundance^11^. Unlike MPRAs, STARR-seq does not rely on synthesis but instead uses fragmented genomic DNA, typically obtained through sonication^15^ or nuclease digestion^16^, enabling genome-wide enhancer screening without sequence-length restrictions^17^. However, STARR-seq also has inherent challenges. The placement of candidate sequences in the 3’UTR can affect mRNA stability, and thereby introduce orientation biases in enhancer quantification^18^. Furthermore, genome-wide STARR-seq requires highly complex libraries, necessitating deep sequencing and high transfection efficiency to achieve sufficient coverage^18–20^. Since random fragmentation rarely generates multiple identical copies of the same fragment, most fragments produce only a single readout. As a result, fragment-level analysis is not feasible, requiring the use of peak-calling algorithm to identify enhancer regions^19,20^. However, these approaches lack the resolution to precisely delineate enhancer boundaries.

In recent years, several MPRA and STARR-seq variants have been developed to facilitate the genome-wide functional characterization of human enhancers and their sequence variants^14,15,21–25^. Among these efforts, the ENCODE Consortium has played a pivotal role by implementing large-scale, high-throughput reporter assays within and across multiple cell lines to systematically map enhancer activity across the genome^26–28^. These efforts have generated extensive datasets that provide a valuable resource for dissecting the regulatory architecture of the genome.

However, several critical questions must be addressed to fully leverage these resources and refine the application of massively parallel reporter assays for deeper functional dissection of enhancer sequences. One key uncertainty is the extent to which the human genome has been functionally characterized. While STARR-seq has the theoretical capacity to screen enhancers genome-wide, practical limitations such as sequencing depth can significantly impact coverage. Additionally, the consistency of enhancer identification across different experimental platforms remains unclear. A recent study systematically compared nine different MPRA and

STARR-seq assay designs using a fixed set of 2,440 sequences, demonstrating how variations in experimental design influence enhancer activity measurements^29^.While this study provided valuable insights, it was conducted under controlled conditions rather than real-world applications, where assay-specific factors—such as library design, sequencing depth, and data-processing pipelines—may introduce systematic biases. Furthermore, the extent to which reporter assays yield consistent regulatory activity profiles and how functionally characterized enhancers align with annotations derived from epigenomic features—such as histone modifications, chromatin accessibility, and transcriptional activity—remain largely unexplored.

To fully integrate these existing large-scale reporter assay datasets for enhancer sequence and functional studies and optimize the future application of massively parallel reporter assays, a systematic evaluation of their genome-wide coverage, cross-assay consistency, and concordance with existing enhancer annotations is needed. Without such an assessment, leveraging these datasets for meaningful biological insights remains challenging, limiting our ability to accurately interpret regulatory landscapes and develop a unified framework for enhancer characterization.

In this study, we systematically evaluated a total of six STARR-seq and MPRA datasets representing four major MPRA and STARR-seq assay types obtained in the human K562 cell line. Initial comparisons of lab-reported enhancer calls revealed limited overlap, prompting a deeper investigation into the factors contributing to cross-assay inconsistencies. We reprocessed all datasets using a unified analytical framework, assessing dataset quality while implementing a standardized enhancer identification pipeline and improving cross-assay comparisons by recording both active and inactive regions. Using this harmonized approach, we found significantly improved enhancer call consistency across assays, especially in cases testing similar sequence composition. Furthermore, we assessed the functional relevance of enhancer candidates defined by enhancer RNA (eRNA) transcription start sites (TSSs) and defined by epigenomic profiles from the ENCODE registry of candidate cis-regulatory elements (cCREs), finding existing enhancer annotations are concordant with massively parallel reporter assay data. We also demonstrated transcription emerged as a critical mark of enhancer function, improving the predictive power of epigenomic features and enhancing the enhancer annotation. Our study provides the first comprehensive assessment of diverse massively parallel reporter assay datasets, offering a framework for integrating these datasets to enhance biological insights and refine functional characterization strategies for future applications.

## Results

### Assessment of Cross-Assay Consistency in Enhancer Identification

We analyzed six distinct STARR-seq and MPRA datasets produced by laboratories within the ENCODE Consortium’s Functional Characterization Center, comprising three TilingMPRA datasets, a LentiMPRA dataset, an ATAC-STARR-seq, and a WHG-STARR-seq dataset^26–28^. Although all assays were performed in the human K562 cell line, they differed in experimental objectives, design strategies, and data processing methods. An overview of these experimental designs is illustrated in Fig. 1a, with detailed dataset descriptions provided in the Supplementary Notes.

**Figure 1.**
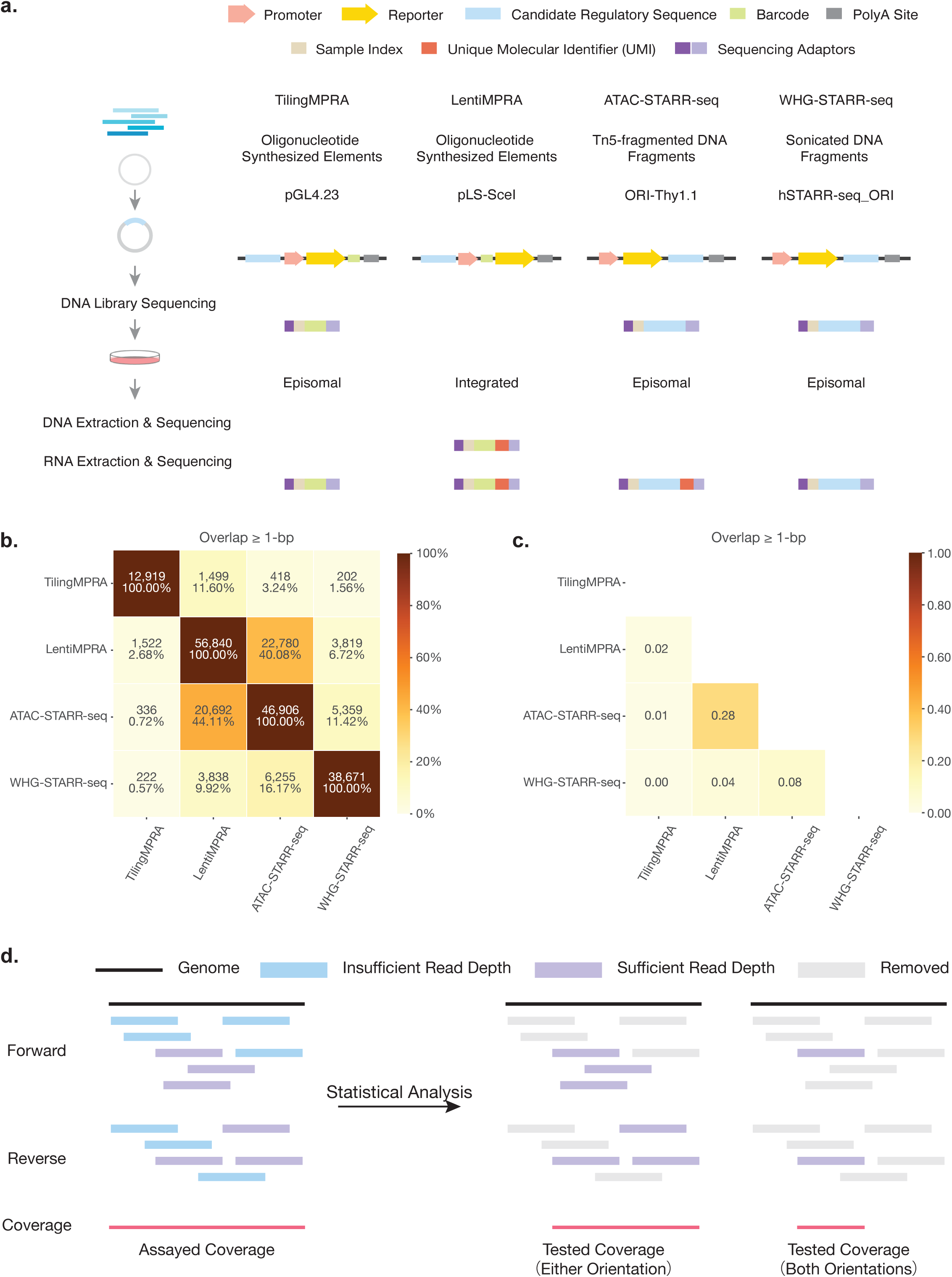
Overview of Experimental Designs and Assay Consistencies Across MPRAs and STARR-seq Assays. **(a)** Schematic representation of the experimental workflows for four types of MPRAs and STARR-seq assays analyzed in this study. **(b)** Heatmap displaying the number of overlapping enhancer regions between assays and their percentage relative to the total number of enhancer regions identified in each assay, based on the ≥1-bp overlap criterion. **(c)** Heatmap presenting the Jaccard Index for pairwise comparisons between assays using ≥1-bp overlap criterion, quantifying overall similarity in enhancer identification. **(d)** Schematic illustration of assayed coverage and tested coverage of reporter assays, distinguishing the proportion of the genome initially assayed versus the regions effectively tested for enhancer activity.

To evaluate the consistency of enhancer identification, we compared enhancer calls reported in each dataset. Data were retrieved from either the ENCODE portal^26–28^ or original publications and processed according to each laboratory’s guidelines. The original number of lab-reported enhancer regions is summarized in Supplementary Table 1. To standardize comparisons, overlapping enhancer calls within each dataset were merged into unique regions, resulting in 12,919 enhancer regions across three TilingMPRA datasets, 56,840 regions from LentiMPRA, 46,906 and 38,671 regions from ATAC-STARR-seq and WHG-STARR-seq, respectively.

We compared enhancer calls across assays by measuring the number of overlapping enhancer regions in each pairwise comparison, applying a minimal overlap threshold of 1 base pair (bp) to ensure inclusion of partially overlapping regions (Extended Data Fig. 1b).The highest overlap was observed between LentiMPRA and ATAC-STARR-seq, where approximately 40% (22,780 out of 56,840) of LentiMPRA regions overlapped with 44% (20,692 out of 46,906) of ATAC-STARR-seq regions. ATAC-STARR-seq and WHG-STARR-seq showed the second-highest overlap, with around 11% (5,359 out of 46,906) of ATAC-STARR-seq regions overlapping with 16% (6,255 out of 38,671) of WHG-STARR-seq regions. Comparisons involving LentiMPRA and WHG-STARR-seq, as well as TilingMPRA with other assays, exhibited lower overlap, reflecting differences in enhancer calls across these datasets (Fig. 1b).

To further quantify similarity across assays, we calculated the Jaccard Index (JI) for each pairwise comparison. Overall, enhancer identification exhibited low consistency, with most JI values approaching zero (Fig. 1c). The highest JI was observed between LentiMPRA and ATAC-STARR-seq (0.28), followed by ATAC-STARR-seq and WHG-STARR-seq (0.08).

Applying stricter overlap criteria further reduced similarities (Extended Data Fig. 2a,b) , highlighting the substantial variability in enhancer identification across different assays.

### Investigating Factors Contributing to Cross-Assay Inconsistencies

The discrepancies in enhancer activity measurements observed across STARR-seq and MPRA assays likely stem from a combination of technical and biological factors. Technical factors include variations in experimental protocols and data analysis methodologies, whereas biological factors encompass aspects such as chromatin context, enhancer-promoter compatibility, sequence positioning, and the inherent properties of the tested sequences.

Chromatin context can affect enhancer activity measurements as episomal assays may not replicate the native chromatin environment, leading to differences in enhancer activities compared to integrated reporter assays, where sequences are chromatinized^23^. Minimal promoter choice in the reporter construct is another source of variability, as different minimal promoters exhibit varying levels of basal transcription^29^. Moreover, certain enhancers respond preferentially to specific promoters, adding further complexity to cross-assay consistency^30–34^.

The positioning of candidate sequences also influences reporter assay outcomes, particularly when characterizing enhancer functions. In many MPRAs, candidate enhancer sequences are placed upstream of a minimal promoter, which may inadvertently measure promoter activity instead of enhancer activity, if the sequence contains promoter-like features^13^. The specific sequences tested also play a substantial role in determining assay outcomes, as each sequence may exhibit distinct regulatory properties and context-dependent behavior. For instance, appending flanking regions to a tested sequence can alter enhancer activity, leading to differences in activity measurements^29^.

Despite the potential impact of these biological factors, a recent study demonstrated that good correlations in enhancer activities can be achieved across different experimental designs in reporter assays—including those used in our study—when a common set of sequences is tested under standardized protocols and analyzed with a unified data processing pipeline^29^. This finding suggests that while biological factors do contribute to variability, they are unlikely to be the primary drivers of the observed inconsistencies. Instead, technical factors, particularly variations in data analysis pipelines, are likely more significant contributors.

A primary factor contributing to the observed cross-assay inconsistencies is the lack of comprehensive reporting of all tested regions. While targeted assays like TilingMPRA and LentiMPRA typically provide quantification data for all tested elements, including both active and inactive elements, genome-wide STARR-seq datasets—such as ATAC-STARR-seq and WHG-STARR-seq—commonly report only the final set of active peak regions. This limited reporting omits regions that were tested but did not reach statistical significance. Without comprehensive information on all regions that proceed through statistical testing, it becomes difficult to determine the true extent of the genome that was functionally evaluated.

Consequently, this limits accurate assessments of genome-wide assay coverage and complicates rigorous cross-assay comparisons. While many current genome-wide STARR-seq analyses assess coverage based on all assayed fragments (Fig. 1d), true assay coverage should reflect only those regions that passed quality filters and underwent statistical testing for enhancer activity, excluding low-coverage regions (Fig. 1d). Additionally, meaningful pairwise comparisons between assays require focusing on regions that were commonly tested across both assays. By reporting only active regions, current comparisons may overestimate inconsistencies, as not all active regions in one assay were necessarily tested in the other. Comprehensive reporting of all tested regions would provide a clearer view of assay coverage and may help reduce the observed inconsistencies, enabling more accurate cross-assay comparisons.

Another important factor contributing to cross-assay inconsistencies is the resolution of enhancer identification, which varies widely between assays and significantly impacts comparisons. High-resolution assays like TilingMPRA and LentiMPRA use fragment-level analysis to define precise enhancer boundaries, while genome-wide STARR-seq typically relies on sliding window peak-calling, generally offering lower resolution. In genome-wide STARR-seq, resolution is heavily influenced by genomic bin size and step size, as these parameters determine the size of the final peak regions. Additionally, methods for calculating read depth in each genomic bin further affect the results. For instance, STARRPeaker^19^, a peak-calling algorithm optimized for genome-wide STARR-seq, has demonstrated that using fragment read depth enables more precise identification of peak summits or centers, aligning with findings that sequence context impacts reporter assay outcomes. Thus, incorporating original fragment boundaries in read depth calculations is essential for consistent and accurate enhancer identification. While some differences in resolution across reporter assays are inevitable, standardizing genomic bin size and step size, and counting only fragments that fully cover each bin into its read depth in genome-wide STARR-seq, should improve cross-assay consistency assessments.

Another factor contributing to cross-assay inconsistencies is orientation bias in enhancer activity measurements, particularly in STARR-seq assays^18^. Although enhancers are generally considered orientation-independent^4^, some assays do not adequately account for orientation in experimental design or data analysis, potentially contributing to observed inconsistencies. For example, TilingMPRA assays tested sequences in only one orientation. In ATAC-STARR-seq and WHG-STARR-seq, although DNA fragments were derived from both orientations, the original analysis pipelines did not separately calculate read depth for genomic bins by orientation. This lack of distinction may have allowed signals from non-overlapping sequences to confound the results, potentially leading to the misclassification of orientation-specific elements as enhancers. Likewise, in LentiMPRA, sequences were tested in both orientations but were classified as enhancers if they were active in either orientation. Without considering activity in both orientations as part of the enhancer-calling criteria, assays may misclassify enhancer regions, complicating cross-assay comparisons. To address this issue, it is essential to test sequences in both orientations and require consistent activity in both orientations as a criterion for enhancer identification. This strategy would improve the accuracy and reliability of enhancer identification and minimize orientation-related biases.

Variations in in-house data processing pipelines may also contribute to the observed inconsistencies across assays. Each pipeline may use distinct probability distributions, bias correction methods, and statistical tests, all of which can affect the results. Additionally, pipelines often apply arbitrary log_2_ fold change thresholds to define active regions, but these thresholds can vary significantly. Given the differences in minimal promoters across assays, normalizing enhancer activity relative to each promoter’s basal transcription level is essential for consistent enhancer identification. Studies suggest that evaluating enhancer activity relative to promoter-specific basal transcription yields more comparable results^35,36^. To improve cross-assay evaluations, implementing a unified approach to process and assess enhancer activity of all datasets is necessary.

### Unified Processing of Reporter Assay Datasets: Initial Data Quality Check

To begin our unified processing of all datasets, we first conducted a comprehensive quality assessment of the reporter assay datasets. In addition to the biological and technical factors discussed previously, the initial quality of these datasets directly impacts assay consistency and reliability.

For TilingMPRA, LentiMPRA, count data were readily available through the ENCODE portal^26–28^, or were re-processed with guidance from the original authors^35^. For ATAC-STARR-seq and WHG-STARR-seq datasets, we applied a unified genomic binning approach, creating 100-bp genomic bins with a 10-bp step size in both forward and reverse orientations (Extended Data Fig. 3a). Only fragments that fully covered each genomic bin were counted, allowing for orientation-independent enhancer identification and a more accurate assessment of genome-wide coverage of tested regions.

We assessed genome-wide coverage for ATAC-STARR-seq and WHG-STARR-seq datasets. Our analysis revealed extensive library complexities of these two datasets, with over 96% of the human genome assayed after processing using the genomic binning approach. However, detailed evaluation identified a notable subset of genomic bins with low read depths in DNA libraries (<10; Methods) (Fig. 2a). This finding raises critical concerns about data quality, likely reflecting limitations in sequencing depth and transfection efficiency. Such limitations suggest that the reported genome-wide coverage of these assays may significantly overstate the regions effectively analyzed, as low-read-depth regions are typically excluded from downstream analyses, thereby reducing the tested coverage of these datasets. Additionally, we evaluated the coverage of accessible regions in both ATAC-STARR-seq and LentiMPRA datasets, given that their assayed fragments were either designed to be enriched in or selected from these regions.

**Figure 2.**
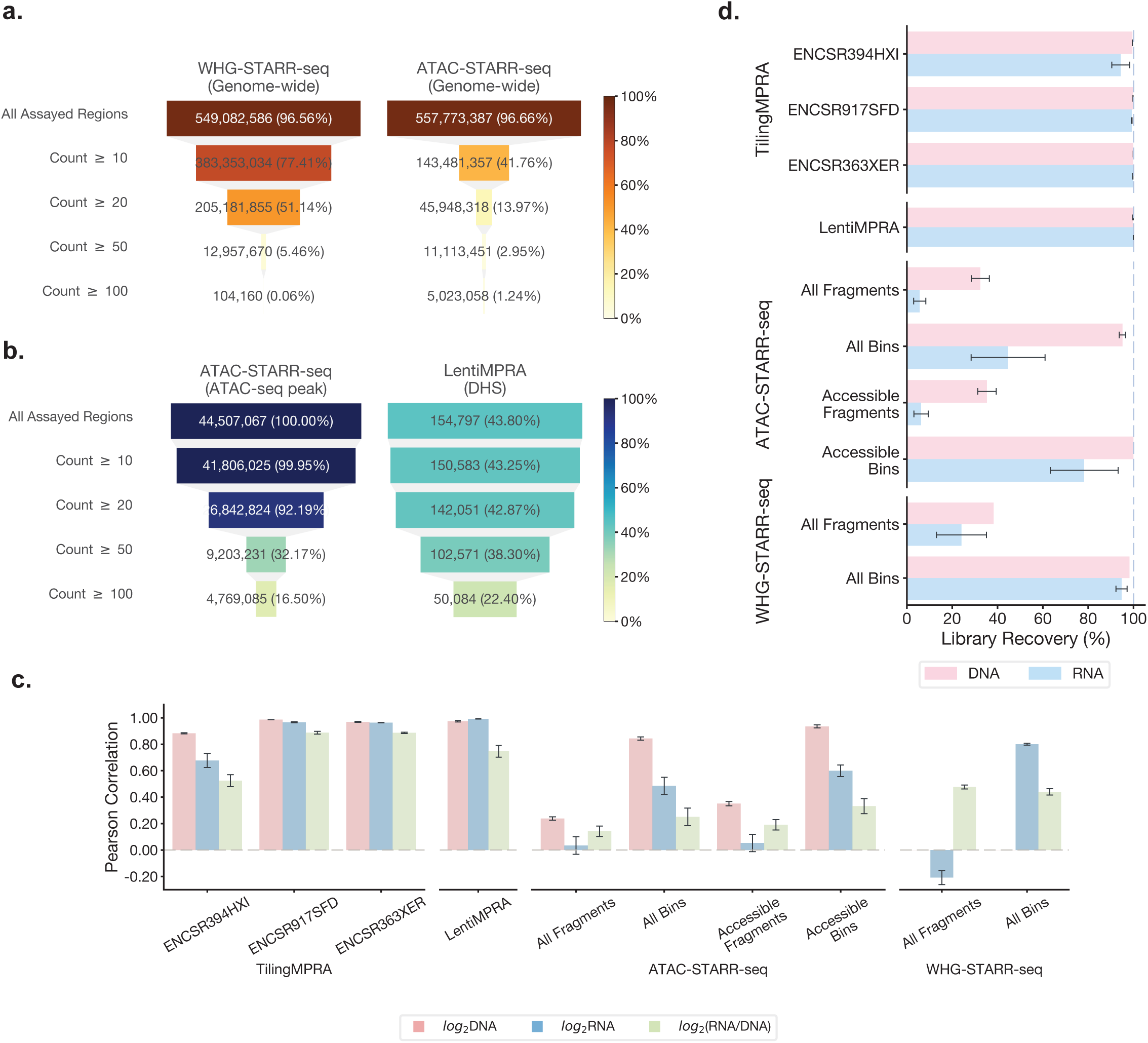
Evaluation of Data Quality Across MPRAs and STARR-seq Assays. **(a)** Funnel plot showing the genome-wide coverage distribution of WHG-STARR-seq and ATAC-STARR-seq at varying read depths thresholds in DNA libraries. **(b)** Funnel plot illustrating the coverage distribution of accessible regions at different read depths thresholds in DNA libraries for ATAC-STARR-seq and LentiMPRA. Accessible regions are defined by ATAC-seq peaks from ATAC-STARR-seq DNA libraries and DNase-seq narrow peaks for LentiMPRA. **(c)** Bar plot presenting average Pearson correlation coefficients for log2-transformed DNA CPM and RNA CPM, and log2(RNA/DNA) ratios across assays. **(d)** Bar plot depicting average library recovery rates in DNA and RNA libraries across assays.

Both datasets demonstrated the ability to capture a substantial proportion of accessible regions with high read depths (Fig. 2b). Specifically, ATAC-STARR-seq achieved almost 100% of coverage of accessible regions characterized by ATAC-seq peaks, while LentiMPRA successfully covered 44% of DNase hypersensitive sites (DHSs) at higher read-depth threshold (≥10) in DNA libraries.

To assess reproducibility between replicates, we calculated Pearson correlations (ρ) for log-transformed counts per million (logCPM) of DNA and RNA counts, as well as log_2_(RNA/DNA) ratios, as these ratios represent the primary measurement of enhancer activity in downstream analyses. Overall, TilingMPRA and LentiMPRA demonstrated strong replicate correlations, indicating high reproducibility across libraries (Fig. 2c). Specifically, LentiMPRA showed robust correlations for both logCPM of DNA and RNA counts (0.97<ρ<0.99) and log_2_(RNA/DNA) ratios (0.72<ρ<0.80). Among the TilingMPRA datasets, ENCSR917SFD and ENCSR363XER displayed consistently high correlations (0.96<ρ<0.99 for logCPM, 0.87<ρ<0.90 for log_2_(RNA/DNA)), while ENCSR394HXI had moderately lower values (0.62<ρ<0.89 for logCPM, 0.47<ρ<0.58 for log_2_(RNA/DNA)), suggesting some variability within this dataset.

In contrast, ATAC-STARR-seq and WHG-STARR-seq demonstrated considerably lower fragment-level reproducibility (Fig. 2c). ATAC-STARR-seq showed weak agreement between replicates (0.001<ρ<0.26 for logCPM, 0.12<ρ<0.22 for log_2_(RNA/DNA)), while WHG-STARR-seq exhibited even greater variability, including negative RNA correlations (Fig. 2c).

Aggregating fragments into genomic bins markedly improved replicate reproducibility for DNA and RNA counts and log_2_(RNA/DNA) ratios in the ATAC-STARR-seq dataset and RNA counts in WHG-STARR-seq dataset (Fig. 2c). Despite these improvements, the correlations for log_2_(RNA/DNA) ratios remained low in both datasets (0.18<ρ<0.37 for ATAC-STARR-seq, 0.42<ρ<0.47 for WHG-STARR-seq. Further restricting analysis to accessible genomic bins in ATAC-STARR-seq provided marginal improvements but did not reach the high reproducibility observed in MPRA datasets, highlighting persistent variability in genome-wide STARR-seq measurements.

We also evaluated library recovery rates by calculating the proportion of fragments or genomic bins with at least one read in each library. TilingMPRA and LentiMPRA had high recovery rates (89%-100%), whereas ATAC-STARR-seq exhibited an average library recovery rate below 40% in DNA libraries and even lower in RNA libraries (Fig. 2d). These findings suggest that many fragments were not consistently detected in ATAC-STARR-seq, possibly due to low sequencing depth or low transfection efficiency. Further analysis of fragments overlapping ATAC-seq peaks showed similar discrepancies in recovery rates between DNA and RNA libraries, pointing to limitations in data quality (Fig. 2d). WHG-STARR-seq also had low recovery rates at the fragment level (18%-38%), but most genomic bins were represented in both DNA and RNA libraries (93%-98%) (Fig. 2d), indicating that issues with sequencing depth and transfection efficiency were not as severe.

These results revealed substantial variability in data quality across different datasets. While MPRA assays exhibited consistently high data quality and reproducibility, genome-wide STARR-seq datasets were more susceptible to limitations such as insufficient sequencing depth and potential low transfection efficiency. These factors likely contributed to higher variability and reduced reliability in enhancer identification, and this issue can remain significant even when genomic binning is applied. Our findings highlight the necessity of applying stringent filtering criteria to exclude low-read-depth regions in the downstream analysis while also ensuring that the final reported tested regions accurately represent sequences with sufficient read depth, rather than using all assayed regions as a proxy for measuring tested region coverage.

### Uniform Processing of Reporter Assay Datasets: Enhancer Call Pipeline

While future studies should further address experimental challenges, to address the role of data processing in contributing to the observed inconsistencies, we implemented a unified enhancer call pipeline and applied it consistently across all datasets. The workflow is illustrated in Fig. 3a, with detailed methodology provided in the Methods section.

**Figure 3.**
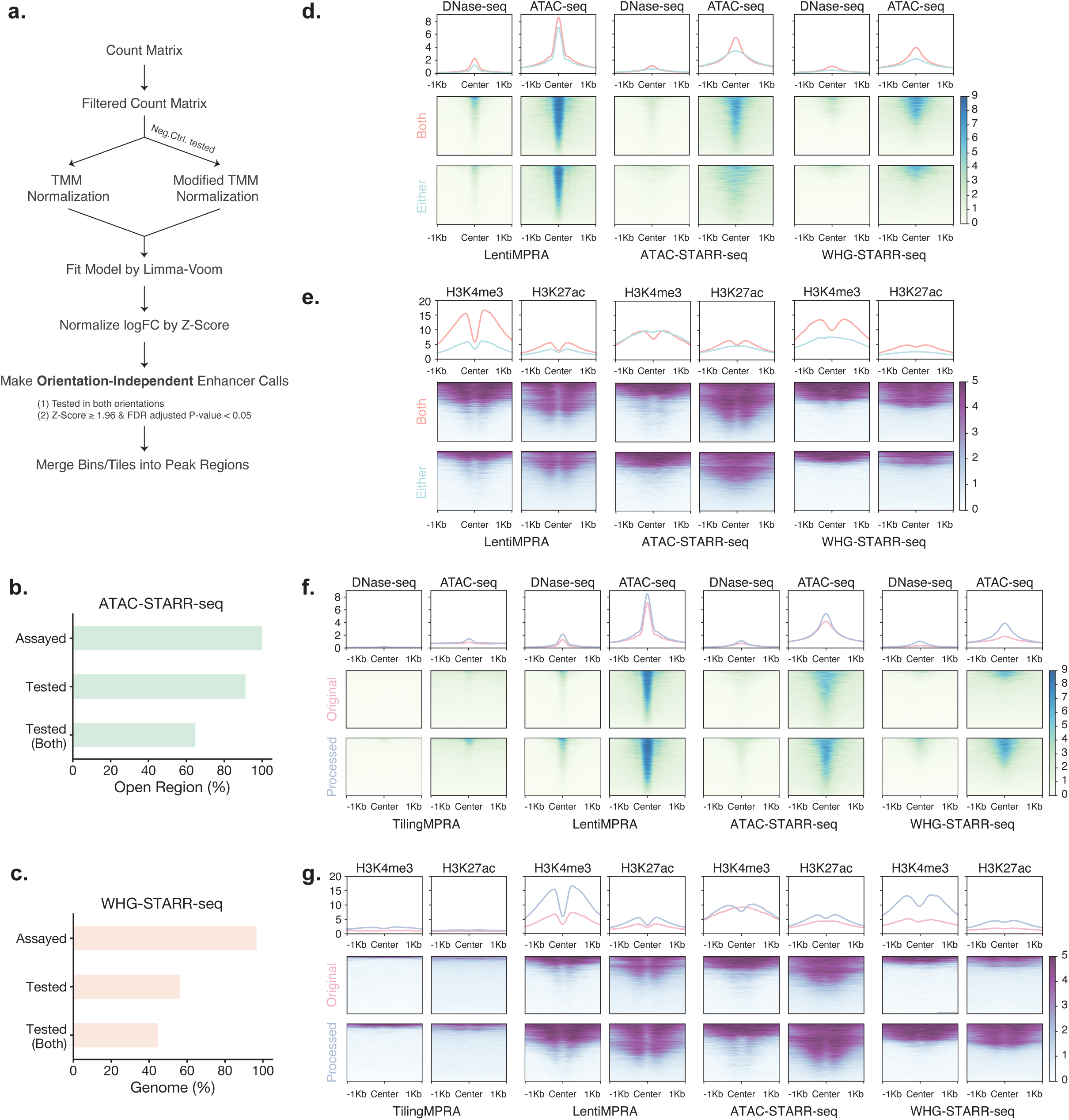
Enhancer Identification Using a Unified Pipeline. **(a)** Schematic of the uniform enhancer call pipeline. The workflow begins with a raw count matrix as input, applies dataset-specific filters to exclude low-depth regions, and normalizes library size using TMM normalization. Regulatory activity is calculated as log2(RNA/DNA) values and Z-score analysis is performed to identify regions with significantly higher regulatory activity than negative control regions as enhancer regions in an orientation-independent manner. **(b)** Bar plot showing the assayed coverage, tested coverage in either orientation and tested coverage in both orientation for open chromatin regions characterized by ATAC-seq peaks derived from DNA libraries in ATAC-STARR-seq. **(c)** Bar plot summarizing genome-wide assayed coverage, tested coverage in either orientation and tested coverage in both orientation. **(d)** Meta-plots comparing the average DNase-seq and ATAC-seq signal profiles (±1 kb from the center) for 2,000 enhancer regions randomly sampled from those identified in both orientations versus 2,000 regions tested in both orientations but active in only one orientation **(e)** Meta-plots comparing the average of H3K4me3 and H3K27ac histone modification profiles (±1 kb from the center) for 2, 000 enhancer regions randomly sampled from those identified in both orientations versus 2,000 regions tested in both orientations but active in only one orientation. **(f)** Meta-plots comparing the average DNase-seq and ATAC-seq signal profile (±1 kb from the center) for 2,000 randomly sampled enhancer regions from laboratory-reported enhancer calls versus those identified using the uniform enhancer call pipeline. **(g)** Meta-plots comparing the average of H3K4me3 and H3K27ac histone profiles (±1 kb from the center) for 2,000 randomly sampled enhancer regions from laboratory-reported enhancer calls versus those identified using the uniform enhancer call pipeline.

The pipeline begins with a raw count matrix as input and applies dataset-specific filters to remove fragments or genomic bins with low read depth. We then adapted the Trimmed Mean of M-values (TMM) normalization^37^ and linear model approach from the Limma-Voom pipeline^38^ to calculate log_2_(RNA/DNA) as a measure of regulatory activity for each fragment or genomic bin in each orientation. For targeted assays that included negative control sequences, we modified the original TMM normalization method to rely solely on negative controls for adjusting library size and composition bias. This approach provides greater accuracy in normalization, particularly for targeted assays where the assumption that most fragments lack regulatory effects may not hold.

After computing the log_2_(RNA/DNA) values, we assessed the regulatory activity of each fragment or genomic bin in both orientations by comparing it to the activity levels of negative controls through a Z-score analysis rather than relying on an arbitrary log2(RNA/DNA) cutoff. This comparison allowed for the identification of regions with significantly elevated activity relative to the basal transcription level defined by the negative controls in each orientation. To mitigate orientation bias, we incorporated regulatory activity in both orientations as a criterion for determining whether a fragment or genomic bin qualifies as a potential enhancer.

For genome-wide STARR-seq datasets that lacked negative controls in the original assays, we used genomic bins within exonic regions as surrogate negative controls, as enhancers are predominantly located in non-coding regions^26,39,40^. To ensure a clear distinction between potential enhancer regions and those likely to exhibit basal transcription, we excluded genomic bins overlapping the 300-bp flanking regions on either side of exons. This approach minimizes the risk of using genomic bins that may have counted fragments overlapping with enhancers in intronic regions, increasing the reliability of these surrogate negative controls.

Finally, our pipeline recorded both active and inactive regions identified in an orientation-independent manner, ensuring an accurate assessment of genome-wide coverage of tested regions. This comprehensive reporting approach also enables robust cross-assay comparisons. Detailed numbers of fragments or genomic bins tested in one or both orientations, the numbers of negative controls, and the numbers of enhancer regions identified are provided in the Supplementary Table 2.

### Improved Enhancer Identification Through Unified Enhancer Call Pipeline

We applied the uniform enhancer call pipeline to all datasets to standardize the identification of enhancer regions. In the ATAC-STARR-seq dataset, while all accessible regions characterized by ATAC-seq peaks were initially included in the assay, 91.20% were statistically tested for regulatory activity in at least one orientation (Fig. 3b). Furthermore, the effective coverage of regions tested in both orientations within accessible chromatin was reduced to 64.72% (Fig. 3b). Similarly, for the WHG-STARR-seq dataset, 96.61% of the entire human genome was included in the assay; however, only 56.15% of regions were statistically assessed in at least one orientation, with just 44.59% tested in both orientations (Fig. 3c). These findings reveal that the effective coverage of genome-wide STARR-seq datasets is significantly lower than expected, underscoring the importance of comprehensive reporting of tested regions to accurately evaluate assay performance and coverage.

Using our unified pipeline, we identified 57 enhancer regions in TilingMPRA (ENCSR394HXI), 16,603 in LentiMPRA, 11,679 in ATAC-STARR-seq, and 25,505 in WHG-STARR-seq.

Notably, these enhancer regions exhibited significant regulatory activity in both orientations. For the two TilingMPRA datasets (ENCSR817SFD and ENCSR363XER), which tested elements exclusively in one orientation, we adapted our pipeline to perform orientation-dependent analysis, identifying 2,117 enhancer regions in ENCSR817SFD and 3,761 in ENCSR363XER.

To evaluate the significance of making orientation-independent enhancer calls, we investigated their epigenomic features by analyzing 2,000-bp windows centered on these regions. Specifically, we compared the epigenomic features of orientation-independent enhancer regions to those of regions that were tested in both orientations but exhibited significant activity in only one, leveraging ENCODE datasets for DNase-seq, ATAC-seq, and ChIP-seq (H3K4me3 and H3K27ac) in the K562 call line. Orientation-independent enhancers displayed higher chromatin accessibility, as indicated by stronger DNase-seq and ATAC-seq signal intensities compared to enhancers active in only one orientation across all datasets (Fig. 3d). Additionally, they exhibited greater enrichment of both promoter- and enhancer-associated histone modifications, with a more pronounced bimodal patter around their centers (Fig. 3e). These findings suggest that orientation-independent enhancers are more robustly marked by epigenomic features characteristic of active regulatory elements and highlight the importance of making orientation-independent enhancer calls.

We also compared the enhancer regions identified through our unified processing pipeline with the original enhancer calls reported by each laboratory. Across all datasets, uniformly processed enhancer regions exhibited higher chromatin accessibility, as evidenced by stronger DNase-seq and ATAC-seq signals (Fig. 3f). Notably, while some enhancer calls from the unified pipeline were in inaccessible regions, they were still more enriched in accessible regions compared to original lab-reported peaks in the WHG-STARR-seq dataset (Fig. 3f). Additionally, histone modification profiles confirmed that orientation-independent enhancer regions identified by the unified pipeline were more strongly marked by H3K4me3 and H3K27ac compared to lab-reported enhancer regions (Fig. 3g). These results highlight the advantages of our unified pipeline in enhancing the confidence of enhancer identification and providing a more reliable foundation for comparative and functional studies.

### Enhanced Consistency Across Assay Using Uniform Processed Enhancer Calls

With both active and inactive regions recorded through our uniform enhancer call pipeline, we reassessed assay consistency by evaluating how many enhancers identified in one assay were also identified as enhancers in others. To achieve this, we conducted pairwise comparisons by assessing the overlap between enhancer regions from one assay and all tested regions in another. Because our enhancer regions were defined in an orientation-independent manner, inactive regions were also generated by merging elements or genomic bins tested in both orientations that lacked significant enhancer activity.

For each pairwise comparison, enhancer regions from assay A were evaluated against all tested regions in assay B, and vice versa, as overlaps were not necessarily symmetric. In cases where an enhancer region overlapped multiple tested regions in another assay, or multiple enhancer regions overlapped a single tested region, we assigned the best overlap based on the highest number of overlapping base pairs to minimize redundancy. We then calculated the JI and recorded both the number of enhancer regions that were also classified as enhancers in the other assay and the total number of enhancer regions tested. By restricting comparisons to commonly tested regions, this approach provided a more accurate and comprehensive assessment of cross-assay consistency.

Using the minimal overlap threshold (≥1-bp overlap), we observed modest improvements in assay consistency, as indicated by higher JI values (Fig. 4a), though the improvement was not statistically significant (one-sided Wilcoxon paired test, *p* = 0.11). However, applying the ≥50% reciprocal overlap threshold resulted in a significant increase in cross-assay consistency, with JI values substantially higher than those based on lab-reported enhancer regions (one-sided Wilcoxon paired test, *p* = 0.02). These findings demonstrate that implementing a uniform enhancer call pipeline and refining comparison strategies improve cross-assay consistency, highlighting the importance of standardized processing in functional characterization studies.

**Figure 4.**
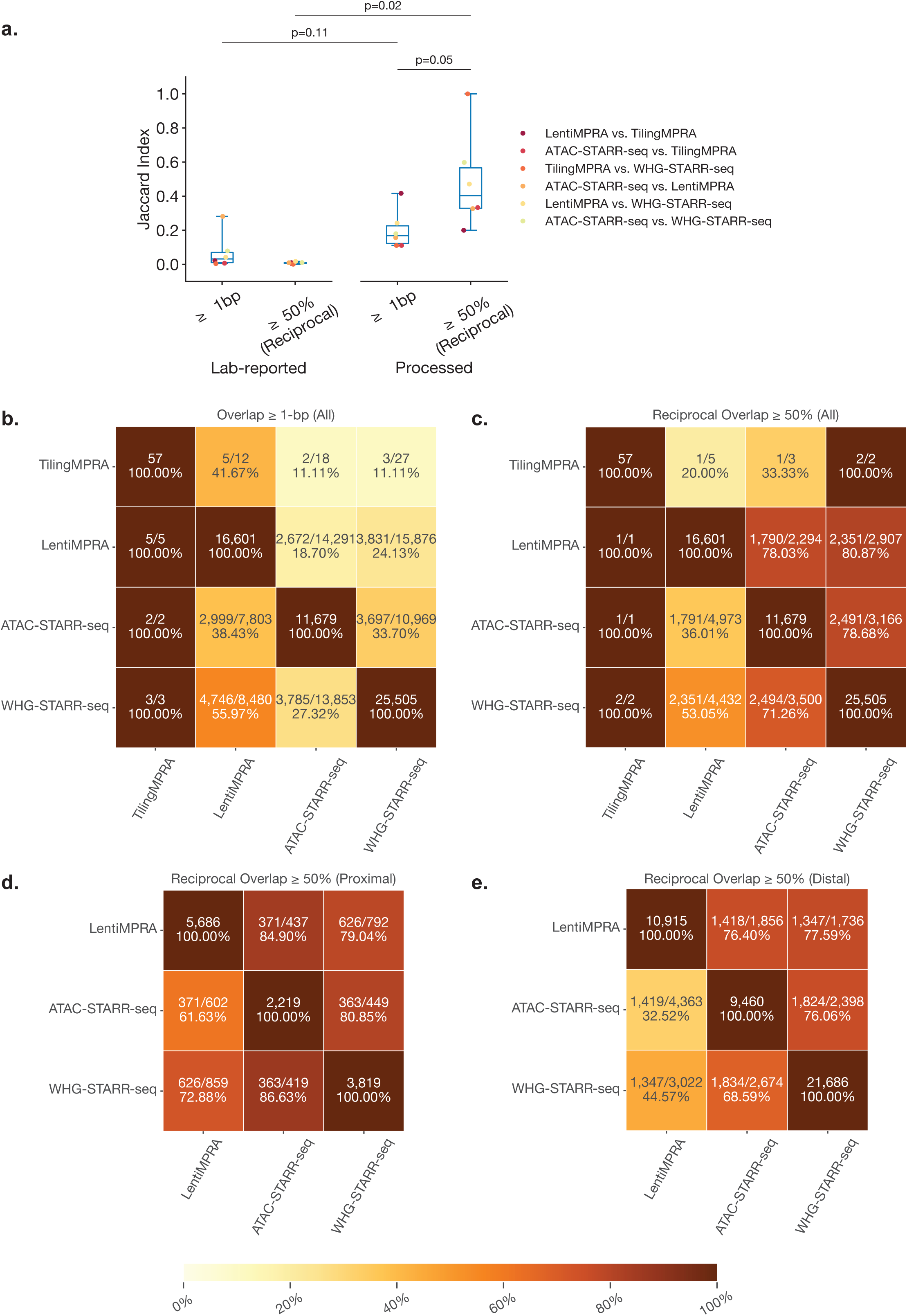
Enhanced Consistency in Cross-Assay Comparisons Using Uniformly Processed Enhancer Calls. **(a)** Box plot showing the Jaccard Index for pairwise comparisons between assays, calculated using the minimal overlap criterion of 1-bp and the stricter criterion of ≥50% reciprocal overlap. Results are shown for both laboratory-reported and uniformly processed enhancer calls, illustrating the improved consistency achieved through uniform processing. **(b,c)** Heatmaps displaying the number of overlapping enhancer regions between assays under the ≥1-bp overlap criterion (b) and under the ≥50% reciprocal overlap criterion (c). Each cell shows the ratio of the number of enhancer regions in the row dataset that overlap with enhancer regions in the column dataset to the number of enhancer regions in the row dataset overlapping with tested regions in the column dataset. Diagonal cells display the total number of enhancer regions identified in each dataset. **(d,e)** Heatmaps displaying the number of overlapping enhancer regions between assays under the ≥50% reciprocal overlap criterion in proximal regions (d) and distal regions (e).

### Sequence Overlap and Assay-Specific Factors Influence Cross-Assay Consistency

While previous comparisons using lab-reported enhancer regions showed lower agreement across assays when a stricter overlap criterion (≥50% reciprocal overlap) was applied, comparisons using uniformly processed data demonstrated the opposite trend: most pairwise comparisons exhibited increased JI values under the stricter criterion compared to the ≥1-bp threshold (Fig. 4a). For instance, when comparing LentiMPRA enhancers to tested regions in ATAC-STARR-seq and WHG-STARR-seq, the proportion of consistently active regions rose from 19% and 24% (using a ≥1-bp threshold) to 78% and 80% (using ≥50% reciprocal overlap), respectively. A similar pattern was observed in pairwise comparisons between ATAC-STARR-seq and WHG-STARR-seq (Fig. 4b,c), indicating that enhancer identification is more consistent when comparing sequences with greater overlap.

Despite the overall increase in consistency with more stringent overlap criteria, assay consistency remained largely unchanged when comparing enhancer regions identified in ATAC-STARR-seq and WHG-STARR-seq to those tested in LentiMPRA, regardless of the overlap threshold (Fig. 4b,c and Extended Data Fig. 4a). This suggests that assay-specific factors, rather than sequence overlap alone, play a dominant role in determining cross-assay agreement for LentiMPRA. Given that LentiMPRA positions candidate sequences upstream of a reporter gene, we suspected that its ability to capture promoter activity rather than enhancer activity is a key factor influencing cross-assay agreement.

To test this, we assessed assay consistency separately in proximal and distal regions. TilingMPRA is excluded from this analysis due to limited sample size. Tested regions were classified as proximal if ≥90% of their sequence overlapped within 500 bp of a protein-coding TSS (based on GENCODE^41^ annotation v45) and distal otherwise. Stratifying comparisons by TSS proximity revealed that ATAC-STARR-seq and WHG-STARR-seq exhibited significantly higher consistency with LentiMPRA in proximal regions than in distal regions. Specifically, ∼62%-73% of proximal enhancer regions identified by STARR-seq assays were also active in LentiMPRA, whereas only ∼33%-47% of distal enhancer regions showed consistent activity (Fig. 4d,e and Extended Data Fig. 4b,c). Notably, these proportions differed only when comparing distal versus proximal regions but remained largely unchanged across different overlap thresholds (Fig. 4d,e). These findings suggest that LentiMPRA is more likely capturing promoter activity rather than enhancer activity as measured in genome-wide STARR-seq assays, emphasizing that assay-specific factors play a dominant role in determining cross-assay consistency when comparing to LentiMPRA.

### Evaluating Functional Support for Enhancer-Like and Promoter-Like Sequences in cCREs

Epigenomic features such as DNA accessibility and histone modifications have long been recognized as key indicators of active enhancers^5,8,40^. Leveraging these features, the ENCODE Consortium established a registry of cCREs^8^. To assess how well these elements are functionally validated by massively parallel reporter assays, we examined their coverage and activity in LentiMPRA, ATAC-STARR-seq, and WHG-STARR-seq datasets.

Since cCREs were not specifically designed as targeted sequences in these assays, we assessed their coverage by identifying overlaps between cCRE elements and tested regions. A cCRE was considered covered if it had at least a 1-bp overlap with a tested region. To further characterize their representation across assays, we categorized covered cCREs into three mutually exclusive groups based on their overlap extent: high (≥80% reciprocal overlap), moderate (50%-80% reciprocal overlap), and low (all other overlap). Detailed coverage statistics are provided in Extended Data Fig. 5a and Supplementary Table 3.

To evaluate the functional relevance of cCREs, we analyzed their active rates across LentiMPRA, ATAC-STARR-seq, and WHG-STARR-seq (Extended Data Fig. 5b). In both genome-wide STARR-seq datasets, cCREs associated with enhancer-like and promoter-like signatures—dELS, pELS, and PLS—demonstrated the highest active rates among all cCRE subtypes, whereas other cCRE categories exhibited lower active rates. Specifically, high-overlap dELS, pELS, and PLS each showed active rates ranging from 46% to 89% in ATAC-STARR-seq and WHG-STARR-seq (Extended Data Fig. 5b), highlighting their strong functional relevance in both genome-wide STARR-seq datasets. In contrast, the active rates of other cCRE subtypes declined sharply, with CA-H3K4me3 and CA-TF elements exhibiting moderate active rates (22%-49%), followed by CA-CTCF and CA-only elements, which showed more limited active rates (5%-9%). As expected, low-DNase elements, which are generally classified as inactive cCREs, displayed the lowest active rates (2%-4%), only slightly higher than regions without any cCRE overlap (0.4%-0.5%).

While the overall active rate patterns were consistent across cCRE subtypes in genome-wide STARR-seq datasets, LentiMPRA exhibited a distinct trend. PLS elements displayed the highest active rate (51%), whereas dELS (18%) and pELS (19%) showed similar activity levels to CA-H3K4me3 (21%), CA-TF (14%), and CA-only (15%) (Extended Data Fig. 5b). These findings again suggest that LentiMPRA may preferentially capture promoter-associated activity rather than enhancer activity, distinguishing it from genome-wide STARR-seq assays.

Additionally, low-DNase elements exhibited an 8% active rate in LentiMPRA, markedly higher than the 2%-4% observed in ATAC-STARR-seq and WHG-STARR-seq. Regions without overlap with any cCREs also showed a higher active rate (2%) compared to the minimal activity levels detected in ATAC-STARR-seq and WHG-STARR-seq (0.4%-0.5%). These results indicate that, beyond its tendency to capture promoter activity, LentiMPRA readouts are influenced by additional assay-specific factors. One possible explanation is that LentiMPRA’s random genomic integration may position low-DNase elements into accessible chromatin regions, artificially increasing their apparent activity.

Collectively, these findings highlight the predictive power of cCREs in identifying active enhancers in reporter assays, particularly for dELS, pELS, and PLS, which exhibited significantly higher activity than other cCRE categories. The near absence of enhancer activity in regions lacking biochemical features underscores the essential role of chromatin accessibility and histone modifications in defining functional enhancers. At the same time, the distinct activity patterns observed in LentiMPRA, likely reflecting its preference for promoter-associated sequences and other assay-specific influences, emphasize the need to carefully consider assay-specific factors when interpreting results and integrating data from different massively parallel reporter assays.

### Transcription as a critical mark of Active Enhancers

In addition to epigenomic features, enhancers are distinguished by their ability to generate eRNAs through divergent transcription^42,43^.Tippens et al. demonstrated that divergent transcription serves as a more precise marker of active enhancers than histone modifications and identified a fundamental enhancer unit based on divergent transcription start sites (TSSs)^18^.

Expanding on this, Yao et al. showed that GRO/PRO-cap is the most effective experimental approach to identify eRNAs and their divergent TSSs, and further compiled an enhancer compendium with a unified definition of enhancers based on divergent transcription^44^.

Leveraging uniformly processed enhancer calls from large-scale reporter assays, we next examined these transcriptional characteristics of enhancers. Using the same analytical framework applied to cCREs, we assessed the coverage of GRO-cap enhancers^44^(divergent elements identified by PINTS from GRO-cap data) across the three assays. Detailed statistics are provided in Supplementary Table 4 and Extended Data Fig. 5d.

High-overlap GRO-cap enhancers exhibited strong enhancer activity, with 87% and 78% being active in ATAC-STARR-seq and WHG-STARR-seq, respectively (Extended Data Fig. 5e). Furthermore, GRO-cap enhancers consistently displayed significantly higher active rates compared to regions that neither overlapped with any GRO-cap elements nor exhibited GRO-cap signals (Extended Data Fig. 5e,f). Notably, while regions devoid of both transcriptional signals and overlap with GRO-cap elements exhibited the lowest active rates across all three assays (0.7%-4%), regions that did not overlap with any annotated GRO-cap elements but still contained detectable GRO-cap signals showed slightly higher, albeit low, levels of activity (2%-11%) (Extended Data Fig. 5f). These findings reinforce the strong functional relevance of GRO-cap enhancers in reporter assays, demonstrating that divergent transcription is a defining characteristic of active enhancers and supporting the enhancer architecture defined by previous studies^18,44^.

To further explore the functional relevance of transcriptional level, we categorized tested regions in LentiMPRA, ATAC-STARR-seq, and WHG-STARR-seq into four transcription-level classes (high, medium, low and none) based on GRO-cap signals^45^ (see Methods) and calculated the active rates within each category. Our analysis revealed that regions with higher transcription levels were significantly more likely to function as enhancers across all three assays (Fig. 5a).

**Figure 5.**
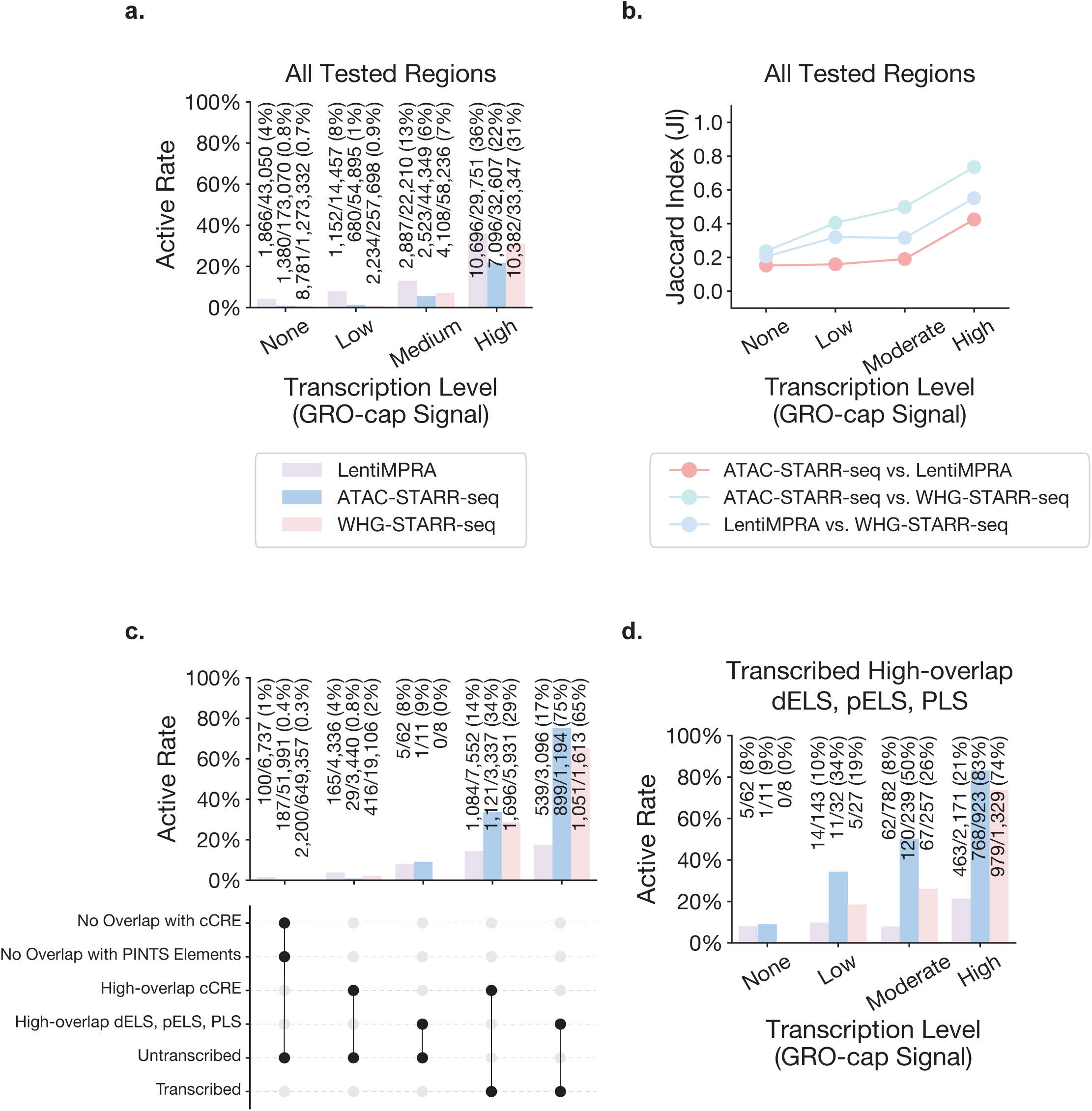
Impact of Transcription Levels on Active Rates and Assay Consistencies. **(a)** Bar plot illustrating the active rates of all tested regions in LentiMPRA, ATAC-STARR-seq, and WHG-STARR-seq, categorized by transcription levels (none, low, medium, and high) determined by GRO-cap signals. (b) Line plot depicting Jaccard Index values for pairwise comparisons between LentiMPRA, ATAC-STARR-seq, and WHG-STARR-seq across all tested regions with varying transcription levels, calculated using the ≥50% reciprocal overlap criterion. (c) Bar plot illustrating the active rate of transcribed and untranscribed regions with high-overlap with any types of cCREs or with high-overlap with dELS, pELS, and PLS or without any overlap with cCRE and PINTS elements. (d) Bar plot showing the active rate of high-overlap dELS, pELS, and PLS regions with different transcription levels (low, medium, high) determined by GRO-cap signals

Regions with no or low GRO-cap signals exhibited minimal enhancer activity, particularly in ATAC-STARR-seq and WHG-STARR-seq, where active rates remained below 1%. Regions with medium transcription level displayed moderate enhancer activity, with active rates around 10% across assays, while highly transcribed regions exhibited the highest active rates, with approximately 30% of tested regions classified as active enhancers (Fig. 5a). These findings reinforce the role of transcription as a key marker of enhancer activity across reporter assays.

Despite the low active rates observed in regions with little or no transcription, thousands of such regions were still identified as active enhancers across all three assays (Fig. 5a). This raised concerns that a subset of these enhancer calls might represent false-positive hits. To explore this possibility, we examined assay consistency across transcription classes, hypothesizing that regions with lower transcription levels would exhibit reduced cross-assay agreement, suggesting a higher prevalence of false positives. Indeed, using ≥50% reciprocal overlap as the comparison criterion, we observed a positive relationship between transcription levels and assay consistency (Fig. 5b, Extended Data Fig. 6). Regions lacking detectable transcription signals exhibited the lowest Jaccard Index values across all pairwise comparisons (Fig. 5b), indicating poor reproducibility across assays. Conversely, high-transcription regions exhibited the highest assay consistencies (Fig. 5b, Extended Data Fig. 6d). These results support the hypothesis that these reporter assays may yield a greater proportion of false positives in regions with lower transcription.

### Transcription Enhances the Predictive Power of Biochemical Features for Enhancer Activity

Next, we assessed whether transcription improves the ability of biochemical features to predict active enhancers. We analyzed tested cCREs with high overlap (≥80% reciprocal overlap) with reporter assay regions and classified them as either transcribed or untranscribed based on detectable GRO-cap signals. We then compared their active rates across assays.

Untranscribed cCREs exhibited low enhancer activity in all three assays (∼0.8%-4%), with active rates only slightly higher than untranscribed regions that lacked cCRE or PINTS annotations (∼0.3%-1%) (Fig. 5c). Untranscribed dELS, pELS, and PLS showed slightly elevated active rates (∼0%-9%), though their sample sizes were limited.

In contrast, transcribed cCREs displayed significantly higher active rates across all assays (∼14%-29%) (Fig. 5c). This trend was particularly pronounced for transcribed dELS, pELS, and PLS, which exhibited much higher active rates (∼17%-75%) than their untranscribed counterparts (Fig. 5c). These results indicate that dELS, pELS, and PLS contain a higher proportion of functional enhancers than other cCRE categories and suggest that transcription serves as an additional predictive layer beyond traditional biochemical features such as chromatin accessibility and histone modifications (H3K4me3 and H3K27ac).

Further stratification of tested dELS, pELS, and PLS by transcription levels reinforced the strong relationship between transcription and enhancer activity across all assay types (Fig. 5d). Highly transcribed dELS, pELS, and PLS exhibited particularly high active rates, reaching 83% in ATAC-STARR-seq and 74% in WHG-STARR-seq (Fig. 5d). These findings emphasize transcription as a critical defining feature of active enhancers, complementing biochemical features and improving the precision of enhancer annotation.

## Discussion

This study provides a comprehensive evaluation of a total of six reporter assay datasets generated by different laboratories, representing four major MPRA and STARR-seq assay types. Our analysis revealed substantial inconsistencies in enhancer identification across assays using original lab-reported enhancer calls, primarily driven by technical variations in experimental workflows and data processing methodologies. By applying a standardized analytical framework, we systematically assessed dataset quality, cross-assay consistency in enhancer identification, and the functional validation of enhancers based on epigenomic features and transcriptional features. Our findings highlight both the strengths and limitations of current high-throughput reporter assays in capturing enhancer activity and underscore the need for standardized experimental and analytical approaches in functional characterization studies.

Through re-processing and quality evaluation of all datasets, we identified insufficient fragment coverage, possibly stemming from inadequate sequencing depth and low transfection efficiency, as the critical limitation in genome-wide STARR-seq assays. These factors compromise not only the reproducibility of enhancer identification but also the effective coverage of tested genomic regions. Particularly in genome-wide assays, large proportions of the genome may remain untested or excluded due to low read depth, leading to an overestimation of genome-wide coverage. Addressing these technical challenges is essential for improving the reliability and completeness of genome-wide enhancer screens.

To address technical discrepancies across assays, we developed and applied a uniform enhancer call pipeline designed to produce orientation-independent enhancer calls. This pipeline incorporated features such as normalization to negative controls, stringent statistical thresholds, and a requirement for enhancer activity in both orientations. Our results demonstrated that this unified approach successfully mitigated many sources of technical variation, yielding a more reliable and consistent set of enhancer regions across datasets. Significantly, our findings emphasized the critical role of orientation-independent analysis and the inclusion of negative controls in enhancing the reliability of enhancer identification. Testing fragments in both orientations and evaluating regulatory activities relative to negative controls proved essential for reducing technical biases. However, we acknowledge that the stringent criteria employed in our pipeline, particularly the requirement for data in both orientations, may have contributed to false-negative results, especially for regions with limited or missing data.

Moreover, the primary goal of our unified enhancer call pipeline was to address technical factors underlying inconsistencies in enhancer identification across assays, rather than to comprehensively optimize sensitivity and specificity for all applications. Future studies should aim to systematically evaluate the trade-offs between sensitivity and specificity in various enhancer call pipelines. Such efforts will be crucial for refining enhancer identification methodologies, particularly as functional characterization assays become increasingly diverse and complex.

Using uniformly processed enhancer calls, we conducted a comprehensive evaluation of cross-assay consistency and found improved agreement in enhancer identification across assays.

Further analysis demonstrated that increasing sequence overlap thresholds substantially improved agreement, particularly in genome-wide STARR-seq datasets. However, LentiMPRA exhibited a distinct pattern, with its enhancer calls showing stronger agreement with STARR-seq assays in proximal regions, reinforcing its tendency to capture promoter-associated activity rather than distal enhancer activity. Additionally, LentiMPRA’s random integration mechanism likely introduces variability by positioning sequences into different chromatin environments, which may either enhance or suppress activity depending on the local chromatin state. These findings emphasize the importance of considering assay-specific characteristics when integrating data from different reporter assays to ensure accurate interpretation of enhancer function.

By evaluating the functional relevance of candidate cis-regulatory elements (cCREs), we confirmed that epigenomic features, such as chromatin accessibility and histone modifications, serve as strong predictors of enhancer activity. cCREs associated with enhancer- and promoter-like signatures—dELS, pELS, and PLS—exhibited significantly higher active rates across genome-wide STARR-seq datasets compared to other cCRE subtypes, reinforcing their biological relevance. Conversely, elements lacking chromatin accessibility and histone modifications displayed minimal activity, underscoring the essential role of these epigenomic features in defining active enhancers.

LentiMPRA, however, displayed distinct activity patterns, with higher active rates for PLS and relatively lower activity for dELS and pELS compared to STARR-seq datasets. These differences suggest that LentiMPRA preferentially identifies promoter-driven regulatory elements rather than enhancers, further highlighting the need to consider assay-specific biases when interpreting MPRA data. Additionally, LentiMPRA showed unexpectedly higher active rates for low-DNase elements, possibly due to its random genomic integration placing these elements into more accessible chromatin regions, altering their apparent activity. These findings reinforce the need for careful interpretation of MPRA data.

Beyond epigenomic features, transcription emerged as a key determinant of enhancer function, with regions with higher transcription level displaying significantly higher activity across reporter assays. High transcription levels were strongly correlated with active rates of tested regions, whereas regions with low or no transcription exhibited greater cross-assay variability, suggesting a higher likelihood of false-positive enhancer calls. This highlights the importance of incorporating transcriptional markers to refine enhancer predictions and reduce misclassification. Furthermore, integrating transcriptional activity with epigenomic evidence improved enhancer annotation, as transcribed cCREs—particularly dELS, pELS, and PLS—showed significantly higher active rates than their untranscribed counterparts. These results suggest that transcription serves as an additional predictive layer beyond traditional chromatin features and should be considered when defining functional enhancers.

This study represents the first systematic evaluation of MPRA and STARR-seq datasets in real-world applications. By identifying critical technical factors and implementing a standardized analytical framework, we provide a foundation for improving experimental protocols and data processing methods in high-throughput reporter assays. Our uniform enhancer call pipeline offers a robust approach to enhancing data consistency and can serves as a benchmark for future studies.

The analytical framework established in this study can be extended to compare results across diverse functional characterization assays, such as CRISPR-based screens. Furthermore, the reliable sets of enhancer regions identified through this pipeline can be leveraged to investigate sequence features, enhancer-promoter interactions, and the structural basis of enhancer activity. Such analyses will deepen our understanding of enhancer biology and elucidate the mechanisms underlying assay-specific variability.

In summary, this study highlights the importance of standardization in enhancer characterization assays and demonstrates the value of integrating transcriptional and biochemical evidence for more accurate enhancer predictions. By addressing the technical and analytical challenges identified here, future studies can advance the functional characterization of human enhancers, ultimately improving our understanding of gene regulation and its implications for human health and disease.

## Methods

### Original Reporter Assay Data Acquisition and Processing

Element quantification data for TilingMPRA datasets and peak regions from ATAC-STARR-seq and WHG-STARR-seq datasets were obtained from the ENCODE portal^26–28^, with corresponding accession numbers listed in Supplementary Table 1. LentiMPRA quantification data and enhancer classifications were retrieved from its original publication^35^ and are also accessible through the ENCODE portal^26–28^.

To define enhancer regions in TilingMPRA datasets, we applied a threshold of log2 fold change (log2FC) ≥ 1 with an adjusted p-value < 0.01. For the ATAC-STARR-seq dataset, regions with log2FC > 0 and an adjusted p-value < 0.01 were classified as enhancer regions. The total number of enhancer regions identified in each dataset, as well as the final numbers after merging overlapping regions, are summarized in Supplementary Table 1.

### Cross-Assay Comparison of Lab-Reported Enhancer Regions

To assess the overlap between enhancer regions reported by different laboratories, we measured the fraction of enhancer regions in one assay that overlapped with enhancer regions identified in another. We performed pairwise comparisons across all datasets using two criteria: a minimal overlap threshold of 1 bp to maximize inclusion of partially overlapping regions (Extended Data Fig. 1b) and a ≥ 50% reciprocal overlap threshold to provide a stricter assessment of enhancer reproducibility (Extended Data Fig. 1c). The number of overlapping enhancer regions was recorded for each pairwise comparison (Fig. 1b and Extended Data Fig. 2a).

### Cross-Assay Comparison of Uniformly Processed Enhancer Regions

To systematically evaluate enhancer identification consistency across assays, we first distinguished enhancer regions from inactive regions in genome-wide STARR-seq datasets. Inactive regions were defined as genomic bins tested in both orientations that did not overlap with any orientation-independent enhancer regions, with overlapping bins merged to form continuous inactive regions.

For each pairwise comparison between assay A and assay B, we first identified orientation-independent enhancer regions in assay A that overlapped with tested regions in both orientations in assay B. The tested regions in assay B included both orientation-independent enhancer regions and inactive regions. We then quantified the proportion of enhancer regions in assay A that were not only tested but also identified as enhancers in assay B. This proportion was calculated as the number of enhancer regions identified in both assays divided by the total number of enhancer regions in assay A that overlapped with tested regions in assay B.

We applied the same two overlap criteria to assess cross-assay consistency: ≥1 bp overlap for broad inclusion and ≥50% reciprocal overlap for a more stringent evaluation. These comparisons were conducted across all datasets, and the results were reported in Heatmaps.

### Jaccard Index Calculations

To quantitatively assess enhancer identification consistency across assays, we computed the Jaccard Index (JI) for each pairwise comparison. The Jaccard Index measures the similarity between two datasets, ranging from 0 to 1, with lower values indicating weaker agreement between assays.

The Jaccard Index for a given pair of assays, A and B, is defined as:

JI(A,B) = |AB||AB|

For comparisons based on lab-reported enhancer regions, A and B represent the sets of enhancer regions identified in two different assays. Given that enhancer regions vary in size across datasets and that multiple enhancer regions in one dataset may overlap multiple regions in another, |AB| is defined as the maximum number of overlapping enhancer regions observed in either direction of comparison (A vs. B and B vs. A). |AB| represents the total number of unique enhancer regions across both assays.

For comparisons based on uniformly processed enhancer calls, A represents the set of orientation-independent enhancer regions in assay A that were also tested in both orientations in assay B, and B represents the corresponding set in assay B tested in both orientations in assay A. |AB| is also determined by the maximum number of overlapping enhancer regions across the two directional comparisons (A vs. B and B vs. A).

### Reprocessing of Genome-Wide STARR-seq Datasets

BAM files for ATAC-STARR-seq and WHG-STARR-seq datasets were retrieved from the ENCODE portal^26–28^. We adapted parts of the STARRPeaker^19^ pipeline to process these BAM files and obtain original fragment counts for each library.

To obtain a refined set of original fragments and their corresponding raw counts, we applied a series of stringent filtering criteria. Unmapped, secondary, and chimeric alignments were discarded to retain only primary alignments. Reads with a mapping quality score below 10 were excluded to ensure high-confidence sequencing data. To mitigate potential biases from PCR amplification, reads with identical genomic coordinates were collapsed, a step applied to DNA replicates in ATAC-STARR-seq and across all WHG-STARR-seq libraries. For RNA libraries in ATAC-STARR-seq, PCR duplicates were removed using unique molecular identifiers (UMIs) to distinguish true biological duplicates from amplification artifacts.

### Genomic Bin Count Generation

To generate genomic bin counts, we used *pybedtools*^46,47^ to partition the human genome into 100-bp bins with a 10-bp step size. For each bin, we summed counts of fragments that fully covered the genomic bin (Extended Data Fig. 3a).

### Quality Assessment of Reporter Assay Datasets: Replicate Reproducibility

To assess the reproducibility of replicates across datasets, we calculated Pearson correlation coefficients (ρ) for log-transformed counts per million (logCPM) of DNA and RNA counts, as well as log2(RNA/DNA) ratios. Correlations were computed between biological replicates within DNA and RNA libraries for each dataset, and the results were averaged to provide an overall measure of replicate reproducibility.

For TilingMPRA and LentiMPRA datasets, replicate reproducibility was evaluated at the fragment level, where enhancer activity was quantified per tested sequence. In contrast, for genome-wide STARR-seq datasets, reproducibility was assessed at both the fragment level and the genomic bin level to account for the different resolution of data processing. Additionally, for the ATAC-STARR-seq dataset, we separately evaluated Pearson correlations in two conditions: across the entire genome and within accessible regions characterized by ATAC-seq peaks identified from DNA libraries in ATAC-STARR-seq.

### Quality Assessment of Reporter Assay Datasets: Library Recovery Rate

The library recovery rate was defined as the proportion of unique fragments detected in a given library relative to the total number of unique fragments identified across the entire dataset, encompassing all DNA and RNA libraries. A fragment was considered part of the dataset’s total unique fragments if it was detected in at least one library, rather than requiring its presence in every library. This total serves as an estimate of the full set of input candidate fragments.

This metric provides insight into the reproducibility of fragment detection across replicates and carries slightly different implications for DNA and RNA libraries. In DNA libraries, higher recovery rates indicate greater consistency in library preparation and sufficient sequencing depth, whereas in RNA libraries, higher recovery rates reflect both efficient transfection and adequate sequencing depth. Conversely, lower DNA library recovery rates may suggest insufficient sequencing depth or stochastic loss of fragments during library preparation, while lower RNA library recovery rates could indicate transfection inefficiencies or suboptimal sequencing depth.

By evaluating library recovery rates for DNA and RNA libraries, we can better assess dataset quality, identifying potential technical limitations affecting dataset quality. The average library recovery rates were calculated for DNA and RNA libraries separately across assays and are presented in Fig. 2d.

### Evaluation of Assay Coverage in the DNA libraries in Genome-wide STARR-seq Datasets

To assess potential limitations in sequencing depth within genome-wide STARR-seq datasets, we examined library complexity and genome-wide coverage by applying various read depth thresholds in DNA libraries. We imposed minimum raw count thresholds of 10, 20, 50, and 100 across all DNA libraries to segment the datasets and evaluate library complexity and genome-wide coverage for each remaining subset of the datasets. These assessments were conducted after binning original fragments into genomic bins.

Since the DNA libraries of genome-wide STARR-seq assays were sequenced before transfection, they reflect the original fragment distribution across the genome. The representation of a region in the input DNA libraries plays a crucial role in determining its likelihood of being transfected and subsequently detected in the output RNA libraries. If a region was underrepresented in the input, it is less likely to have been thoroughly tested for enhancer activity in the RNA output.

We utilized *pybedtools*^46,47^ to quantify the genomic coverage by computing the number of base pairs covered at each threshold. The percentage of genome-wide coverage was determined by dividing the number of covered base pairs by the total number of base pairs in the hg38 human reference genome^48^.

For ATAC-STARR-seq, which was specifically designed to enrich open chromatin regions rather than provide full genome coverage, we additionally evaluated its coverage within open chromatin regions, as defined by ATAC-seq peaks from its DNA libraries. The open chromatin coverage of ATAC-STARR-seq was calculated by dividing the number of base pairs covered within ATAC-seq peaks by the total number of base pairs in these peaks.

To provide a comparison, we also assessed open chromatin coverage for LentiMPRA, as its candidate sequences were selected from DNase-seq peaks. DNase-seq peak regions were obtained from the ENCODE portal^26–28^ (Accession: ENCFF185XRG). The open chromatin coverage for LentiMPRA was calculated using the same approach as in ATAC-STARR-seq, by determining the fraction of base pairs covered within DNase-seq peak regions.

### Uniform Enhancer Call Pipeline

To quantify enhancer activity based on log₂(RNA/DNA) ratios, we adapted the Limma-Voom pipeline^49^ with key modifications tailored to different datasets. While the linear model framework was retained, we implemented dataset-specific filtering strategies, a modified TMM normalization approach, and a Z-score-based classification method to identify enhancer regions in an orientation-independent manner.

### Uniform Enhancer Call Pipeline: Dataset-Specific Filtering Strategy

Our filtering strategy was adapted from the filterByExpr function in edgeR^50^. Initially, raw counts were transformed into log-counts per million (logCPM) to normalize for variations in library size. The filtering threshold was determined by computing the logCPM equivalent of a predefined raw count cutoff, ensuring sufficient read depth for reliable downstream statistical analysis.

Fragments were retained if their logCPM values exceeded the threshold across all DNA libraries or, in cases where stricter filtering would excessively reduce coverage, in a minimum required number of libraries, regardless of whether they were DNA or RNA libraries. This approach ensured a balance between stringent filtering for reliable enhancer activity detection and preserving sufficient genome-wide coverage across diverse reporter assay datasets.

For TilingMPRA and LentiMPRA, the filtering threshold was determined using the smallest DNA library, applying a raw count threshold of 10 to establish the corresponding logCPM cutoff. In ATAC-STARR-seq, to accommodate the larger complexity of the dataset while maintaining sufficient coverage, the threshold was calculated using the average DNA library size with a raw count of 20. In WHG-STARR-seq, where only a single DNA library was available, the logCPM threshold was instead based on the smallest library across both DNA and RNA libraries, using a raw count of 10. Unlike the other assays, where filtering was applied across all DNA libraries, WHG-STARR-seq employed a more lenient criterion, retaining fragments if they met the logCPM threshold in at least two libraries, regardless of type. This dataset-specific adaptation ensured that filtering remained stringent enough to remove fragments with low read depth while preserving sufficient genome-wide coverage for reliable enhancer identification.

### Uniform Enhancer Call Pipeline: Normalization Strategy

To normalize for library size differences, we applied the Trimmed Mean of M-values (TMM) normalization from edgeR^37,50^, following the standard Limma-Voom framework. In ATAC-STARR-seq and WHG-STARR-seq, where most genomic bins were expected to exhibit no regulatory activity (i.e., showing no significant difference between RNA and DNA libraries), we applied the conventional TMM normalization method, assuming that the majority of regions had minimal transcriptional changes.

For LentiMPRA and TilingMPRA, we implemented a modified TMM normalization approach to address assay-specific biases. These assays included designated negative control sequences, and in the case of LentiMPRA, candidate sequences were particularly enriched for protein-coding promoters and potential enhancer elements^35^. This enrichment could result in a dataset disproportionately composed of regulatory-active fragments, making the standard assumption that most fragments were not differentially expressed less applicable. To address this, we modified TMM normalization to rely exclusively on negative control elements, allowing for a more accurate adjustment of library size and composition biases without being influenced by the overrepresentation of active regulatory elements. This refinement optimized normalization for the unique design of these assays, ensuring more reliable quantification of enhancer activity.

### Uniform Enhancer Call Pipeline: Enhancer Classification and Statistical Significance

Using the Limma-Voom procedure, log2(RNA/DNA) ratios were then calculated to quantify enhancer activity for each fragment or genomic bin. Rather than applying an arbitrary threshold, we employed a Z-score-based approach to identify regions with significantly elevated activity compared to background transcription levels. Background transcription levels were estimated using negative control elements, and the log2(RNA/DNA) threshold was set at the 95th percentile of negative control distributions.

For genome-wide STARR-seq datasets that lacked dedicated negative controls, genomic bins located within exonic regions were used as surrogate controls to determine the log2(RNA/DNA) threshold. This approach was based on previous studies indicating that enhancers are primarily located in non-coding regions^26,39,40^. To further refine enhancer identification and mitigate orientation bias, enhancer regions were required to exhibit significant activity in both forward and reverse orientations. First, fragments or genomic bins tested in both orientations were identified. Regions were then classified as enhancers if their log2(RNA/DNA) ratios exceeded the threshold and had an adjusted p-value < 0.05. Overlapping fragments or genomic bins were merged to generate the final set of enhancer regions, ensuring a robust and unbiased identification process.

### Uniform Enhancer Call Pipeline: Comprehensive Reporting and Dataset Summary

The pipeline provided a comprehensive reporting of both active and inactive regions, ensuring accurate estimation of genome-wide coverage and facilitating robust cross-assay comparisons.

For genome-wide STARR-seq datasets, we reported multiple levels of coverage to reflect the extent of assay representation. *Assayed coverage* included all genomic bins before applying filters, representing the initial set of regions targeted in the experiment. *Tested coverage* encompassed genomic bins that remained after applying filtering criteria, reflecting regions with sufficient read depth for reliable enhancer activity quantification. Additionally, *tested coverage in both orientations* was defined as the subset of tested genomic bins that were assayed in both forward and reverse orientations, ensuring a stringent assessment of enhancer activity independent of strand bias.

The final dataset included log2(RNA/DNA) ratios, Z-scores, and statistical significance metrics for all tested genomic bins, as well as for bins tested in both orientations. Additionally, merged orientation-independent enhancer regions were reported to provide a set of enhancer calls across datasets. A summary of enhancer region counts, negative controls, and tested regions for each dataset is provided in Supplementary Table 2.

### Evaluation of Genomic Context to Compare Regions

To examine the genomic context of enhancer regions, including DNA accessibility and histone modifications (H3K4me3 and H3K27ac), we compared lab-reported enhancer regions with uniformly processed enhancer regions and orientation-independent enhancers with regions tested in both orientations but identified as active in only one orientation. We utilized publicly available datasets from the ENCODE portal^26–28^, specifically DNase-seq data (ENCFF972GVB), ATAC-seq data (ENCFF102ARJ), H3K4me3 ChIP-seq data (ENCFF911JVK), and H3K27ac ChIP-seq data (ENCFF381NDD).

For visualization, we used deepTools^48^ to generate metaplots of signal intensities across enhancer regions. To compare lab-reported enhancer regions with uniformly processed enhancer regions, we randomly sampled 5,000 enhancer regions from each dataset. Signal intensities were plotted within a 2,000-bp window centered at the region midpoint to capture local epigenomics features.

For the comparison between orientation-independent enhancer regions and those tested in both orientations but were active in only one orientation, we first selected genomic bins that were tested in both orientations. We then identified bins that were active in only one orientation, merged overlapping bins into contiguous regions, and excluded any regions that overlapped with orientation-independent enhancers. From each dataset, we randomly sampled 5,000 orientation-independent enhancer regions and 5,000 regions that were active in only one orientation for comparative analysis.

### Negative Control Regions in Genome-wide STARR-seq Datasets

Since the original ATAC-STARR-seq and WHG-STARR-seq assays did not include dedicated negative controls, we leveraged their genome-wide coverage to define genomic bins overlapping exonic regions as surrogate negative controls. This allowed for the implementation of a Z-score approach in these datasets to establish a background transcription level for enhancer classification.

To identify suitable genomic bins as negative controls, we extracted all exons of protein-coding genes from the GENCODE v42 annotation (hg38)^51^. Because the original fragments covering the 100-bp genomic bins could be substantially longer, we further excluded 300-bp flanking regions on both sides of each exon to prevent potential overlap with adjacent intronic regions, which could confound the regulatory activity measurements of exonic genomic bins. This filtering step ensured that only mid-exonic regions were retained as the final negative control reference regions.

We then identified all genomic bins that were fully contained within these negative control reference regions. These bins were used exclusively in the Z-score approach to characterize background transcription levels but were not used in the TMM normalization process.

### Analysis of Coverage and Active Rate of cCREs and GRO-cap Enhancers

The comprehensive reporting of both active and inactive regions in genome-wide STARR-seq datasets enabled a systematic evaluation of the coverage and active rates of alternative enhancer annotations, such as cCREs and GRO-cap enhancers, across assays.

We retrieved cCRE annotations for K562 cells (Accession: ENCFF286VQG) from the ENCODE Portal^26–28^. GRO-cap enhancers were defined as divergent elements identified by PINTS^44^.

To determine the extent to which these enhancer annotations were tested in LentiMPRA, ATAC-STARR-seq, and WHG-STARR-seq, we applied three mutually exclusive overlap categories: (1) high-overlap (≥80% reciprocal overlap), (2) moderate-overlap (50%-80% reciprocal overlap), and (3) low-overlap (all other overlaps). The number of cCREs and GRO-cap enhancers tested in each assay is provided in Supplementary Table 3 and Supplementary Table 4.

To assess the functional relevance of these tested elements, we calculated their active rates within each overlap category. The active rate for each element type was defined as the proportion of tested elements that exhibited significant regulatory activity.

### Annotation of Transcription Levels for Tested Regions Using GRO-cap Signals

To compare the active rates of tested regions with different transcription levels, we annotated each tested region with transcription levels based on GRO-cap signal data extracted from bigWig files^45^.

To quantify the transcriptional activity within each tested region, we summed the GRO-cap signal from both orientations and normalized it by the region size. The normalized transcription level for each region was computed as the total GRO-cap signal divided by the length of the tested region.

Based on the normalized GRO-cap signal, we classified transcription levels into four categories: (1) None, for regions with no detectable GRO-cap signal; (2) Low, for regions with normalized GRO-cap signal ≤ 0.01; (3) Medium, for regions with normalized GRO-cap signal > 0.01 and ≤ 0.08; and (4) High, for regions with normalized GRO-cap signal > 0.08.

## Supplementary Information

### MPRA and STARR-seq Datasets Utilized in This Study TilingMPRA

This study includes three TilingMPRA datasets generated by the same laboratory, designed to screen enhancers within selected gene loci using a tiling approach with overlapping 200-bp sequences.The first dataset, ENCSR394HXI, assayed tiling sequences in both orientations with a 5-bp sliding window across the *FEN1, FADS1, FADS2,* and *FADS3* loci. The second dataset, ENCSR917SFD, tested tiling sequences in the forward orientation with a 50-bp sliding window at the *MYC* and *GATA1* loci. The third dataset, ENCSR363XER, analyzed tiling sequences in the forward orientation using a 100-bp sliding window, targeting the *LMO2, HBE1, RBM38, HBA2,* and *BCL11A* loci.

The experimental design of these TilingMPRAs follows the classic MPRA framework, utilizing the pGL4.23 vector, with barcodes incorporated into the 3′ UTR of the reporter gene. These assays were performed episomally. Negative and positive controls were included in all three datasets, and candidate elements were synthesized using oligonucleotide synthesis. DNA libraries were sequenced prior to transfection, with stringent control of PCR cycles to minimize amplification bias. The datasets were originally analyzed using DESeq2^52^ compute log₂ fold changes between RNA and DNA libraries and assess statistical significance. Normalization was applied relative to the distribution of negative controls, and enhancer activity was assigned to elements with a log₂ fold change ≥ 1 and an adjusted *p*-value < 0.01.

### LentiMPRA

The LentiMPRA dataset was designed to characterize putative enhancers and promoters selected from DNase I hypersensitive sites (DHSs) and tiling sequences outside DHS regions. These sequences were chosen from loci including *GATA1, MYC, HBE1, LMO2, RBM38, HBA2,* and *BCL11A,* with additional positive and negative controls^29,35^. The 200-bp elements were synthesized via oligonucleotide synthesis.

LentiMPRA utilizes the pLG-Scel vector, which integrates candidate sequences into the genome via lentiviral delivery, with barcodes located in the 5′ UTR of the reporter gene^35^. Unlike TilingMPRA, both DNA and RNA libraries were collected from cells post-integration. To distinguish biological duplicates from PCR duplicates, unique molecular identifiers (UMIs) were included in the sequencing library before sequencing. The dataset was processed using MPRAflow^35,53^ computing log₂ fold changes per element across replicates. Normalization was performed within each replicate, and the final log₂ fold change for each element was calculated as the average across all replicates. Elements with log₂ fold changes exceeding the 95th percentile of negative controls were classified as enhancers^35^.

### ATAC-STARR-seq

ATAC-STARR-seq integrates ATAC-seq (Assay for Transposase-Accessible Chromatin with high-throughput sequencing) with STARR-seq to identify enhancers active in accessible chromatin regions. By leveraging the open chromatin landscape, this approach facilitates the functional validation of active enhancers.

The assay utilized the ORI-Thy1.1 vector in an episomal context, allowing enhancer activity to be measured independently of chromatin context effects. DNA libraries were sequenced prior to transfection, and UMIs were incorporated exclusively into RNA libraries. PCR duplicates were removed from DNA libraries by collapsing fragments with identical genomic coordinates.

ATAC-STARR-seq data were analyzed using CSAW (ChIP-Seq Analysis with Sliding Windows)^54,55^. This method employs a genome-wide sliding window approach, quantifying fragment overlaps and identifying differential activity between RNA and DNA libraries via the quasi-likelihood framework in edgeR^54–56^. Enhancer activity was assigned to genomic regions exhibiting transcriptional activity (*log₂ fold change ≥ 0*) and statistically significant enrichment (*adjusted p-value < 0.05*).

### WHG-STARR-seq

WHG-STARR-seq was designed for genome-wide enhancer identification using randomly fragmented DNA^15^. The DNA libraries were sequenced before transfection to establish baseline representation.

The assay employed the hSTARR-seq_ORI vector. Unlike the targeted approaches of LentiMPRA and TilingMPRA, WHG-STARR-seq screened fragmented genomic sequences without predefined selection criteria. Data were processed using CRADLE^20^ (Correction of Read Counts and Detection of Locally Enriched Regions), which corrects for sequence-based read count biases and identifies significantly enriched regions based on base-pair-level read density.

## Acknowledgements

This work was supported by the National Institutes of Health grants UM1HG009393, R01 HG012970, and R01 AG077899 to J.T.L. and H.Y., as well as R01 DK127778 and R01 DK115398 to H.Y. We thank members of the Yu and Lis laboratories for insightful discussions and feedback throughout this study. We are also grateful to the ENCODE Consortium, particularly T.E. Reddy, K. Siklenka, A. Barrera, Y. Kim, R. Tewhey, H. Dewey, V. Agarwal, A. Du, and D. Lee, for their valuable discussions and guidance.

## Author Contributions

J.Z. conceptualized the study with guidance from H.Y., J.T.L., N.T., and J.L. J.Z. developed the methodologies, performed the formal analysis, and generated the figures. L.Y., A.L., Y.Z., A.W., X.P., K.S., A.B. and T.E.R. contributed to data curation and testing the pipeline code. J.Z. wrote the manuscript with guidance from H.Y. and J.T.L. A.O., Z.Z., L.Y., and K.S. reviewed and edited the manuscript. H.Y. and J.T.L. oversaw the project and acquired funding.

## Ethics Declarations

The authors declare that there are no competing interests.

---

**Extended Data Figure 1.**
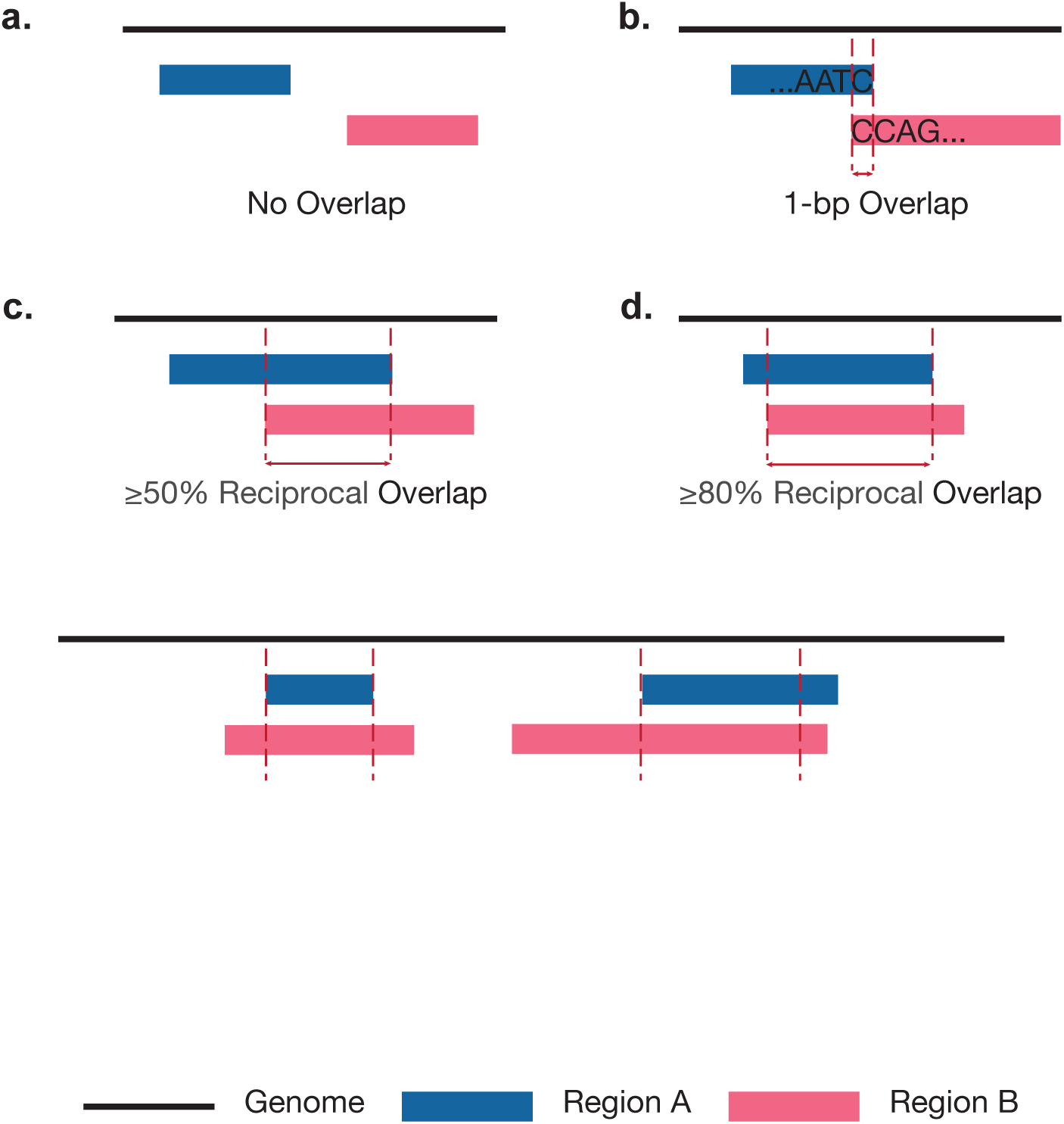
Schematic Representation of Overlap Criteria. **(a)** No overlap between regions A and B. (b) 1-bp overlap between regions A and B. (c) ≥50% reciprocal overlap between regions A and B, where both regions share at least 50% of their length with one another. (d) ≥80% reciprocal overlap between regions A and B, where both regions share at least 80% of their length with one another.

**Extended Data Figure 2.**
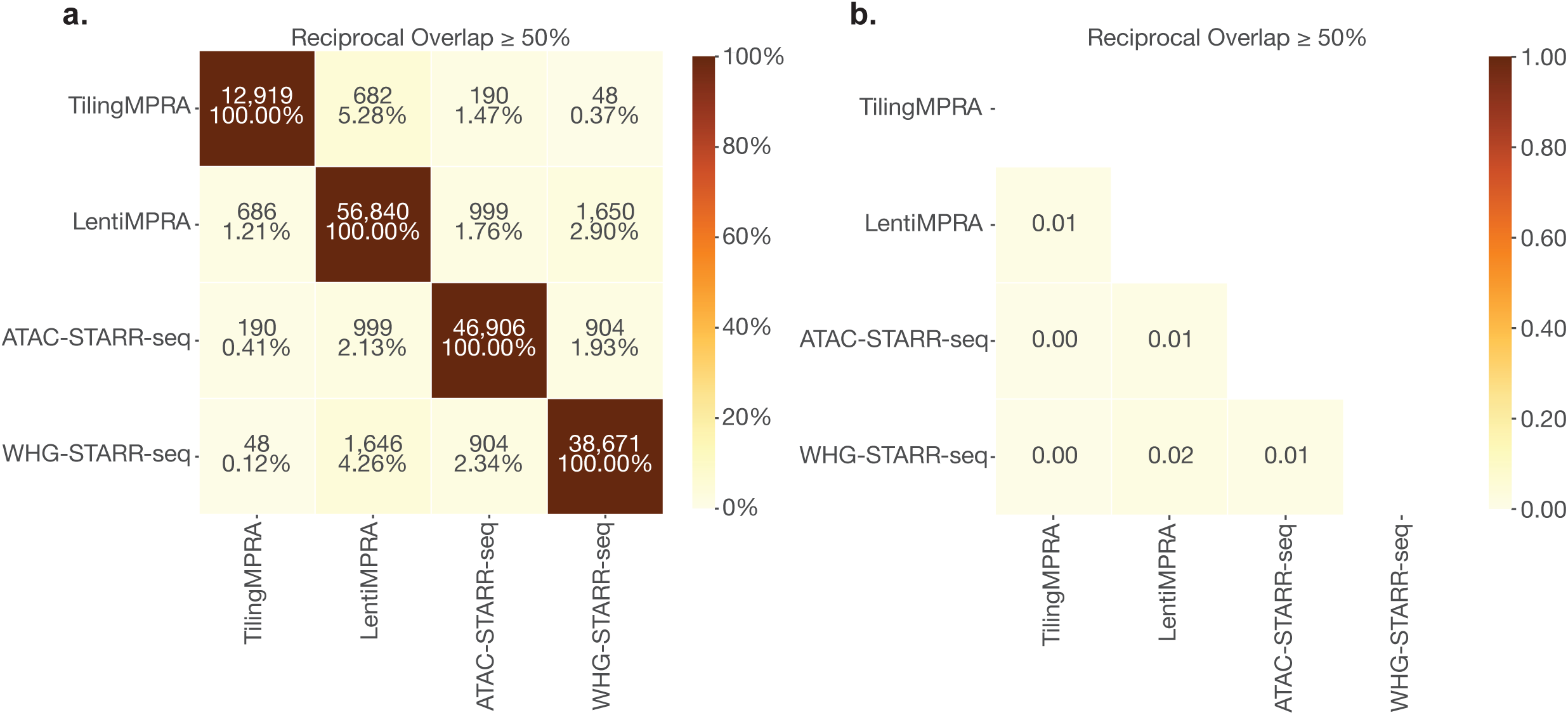
Assay Consistency in Cross-Assay Comparisons Using Original Laboratory-Reported Enhancer Calls. **(a)** Heatmaps showing the number of overlapping enhancer regions between assays under the ≥50% reciprocal overlap criterion. Each cell displays the number of enhancer regions in the row dataset overlapping with those in the column dataset, with diagonal cells indicating the total number of enhancer regions identified in each dataset. **(b)** Heatmap presenting the Jaccard Index of each pairwise comparison between assays using the ≥50% reciprocal overlap criterion.

**Extended Data Figure 3.**
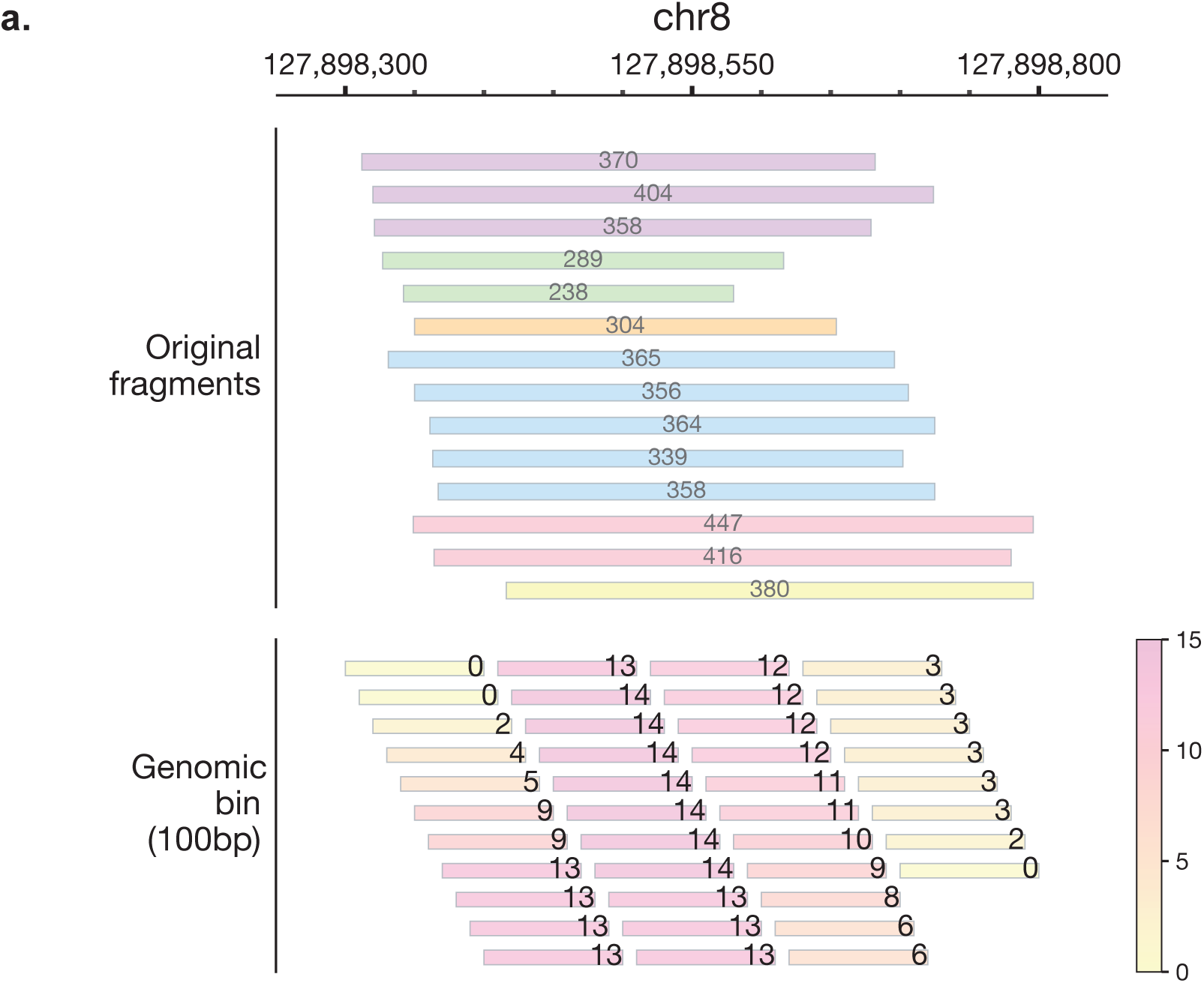
Schematic Representation of Binning Strategies in Genome-wide STARR-seq Datasets. **(a)** Illustration of the genomic binning approach applied to ATAC-STARR-seq and WHG-STARR-seq datasets. Original STARR-seq fragments are assigned to 100-bp genomic bins using a 10-bp sliding window, where only fragments fully covering the genomic bin are counted.

**Extended Data Figure 4.**
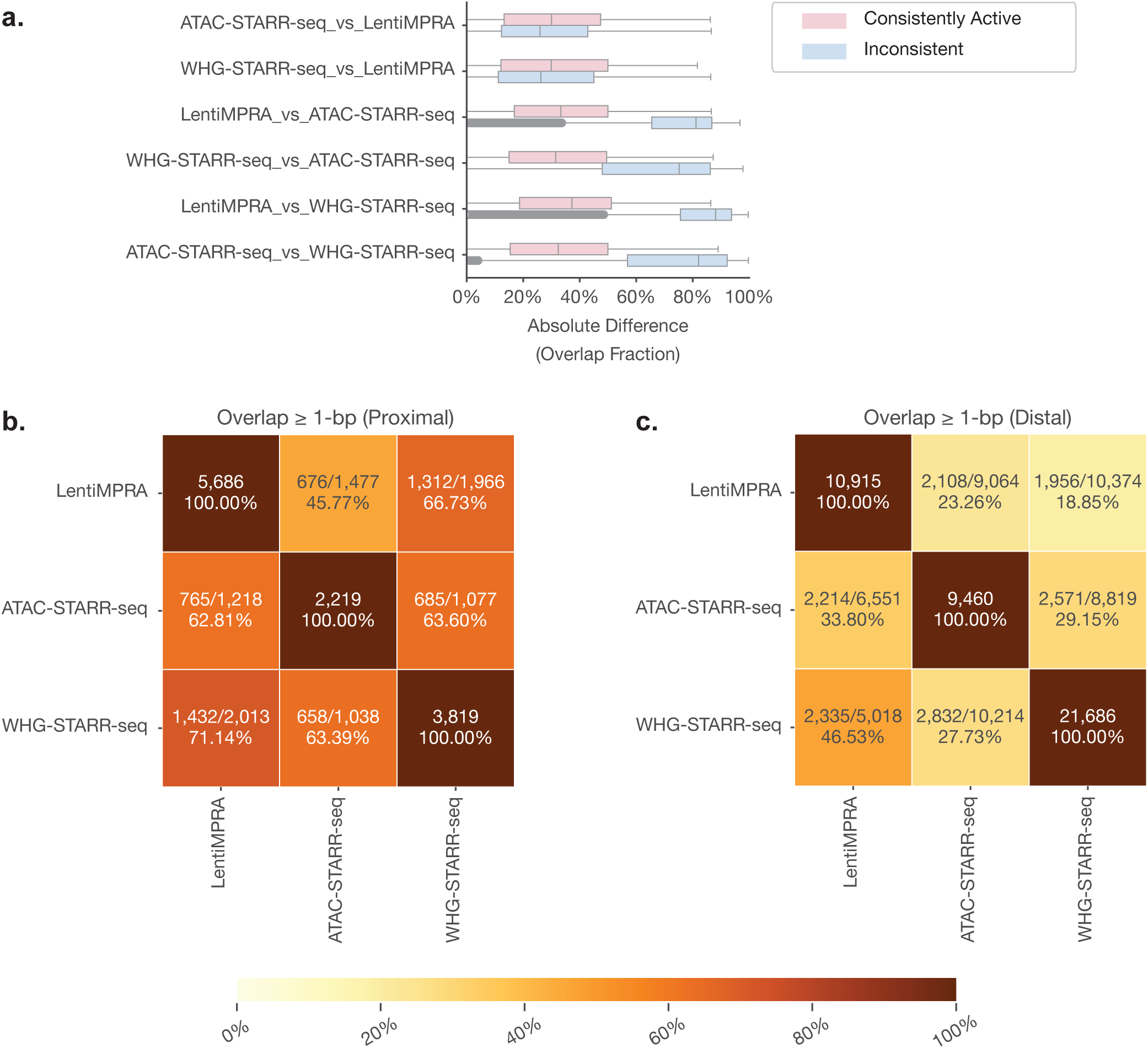
Assay Consistencies for Distal and Proximal Regions Tested Across MPRA/STARR-seq Assays. **(a)** Box plot illustrating the absolute difference in overlap fraction for each pairwise comparison between assays, distinguishing consistently active regions from those with inconsistent calls. The absolute difference in overlap fraction quantifies sequence similarity based on relative overlap proportions, calculated as the number of overlapping base pairs divided by the total region length in assay A. The absolute difference between these values reflects variations in sequence length, where lower values indicate greater sequence similarity, while higher values suggest larger differences in sequence composition. **(b, c)** Heatmaps displaying the number of overlapping tested regions between assays under the ≥1-bp overlap criterion in proximal regions (b) and in distal regions (c). Each cell shows the ratio of the number of proximal enhancer regions in the row dataset that overlap with proximal enhancer regions in the column dataset to the number of proximal enhancer regions in the row dataset overlapping with proximal tested regions in the column dataset. Diagonal cells display the total number of proximal enhancer regions identified in each dataset.

**Extended Data Figure 5.**
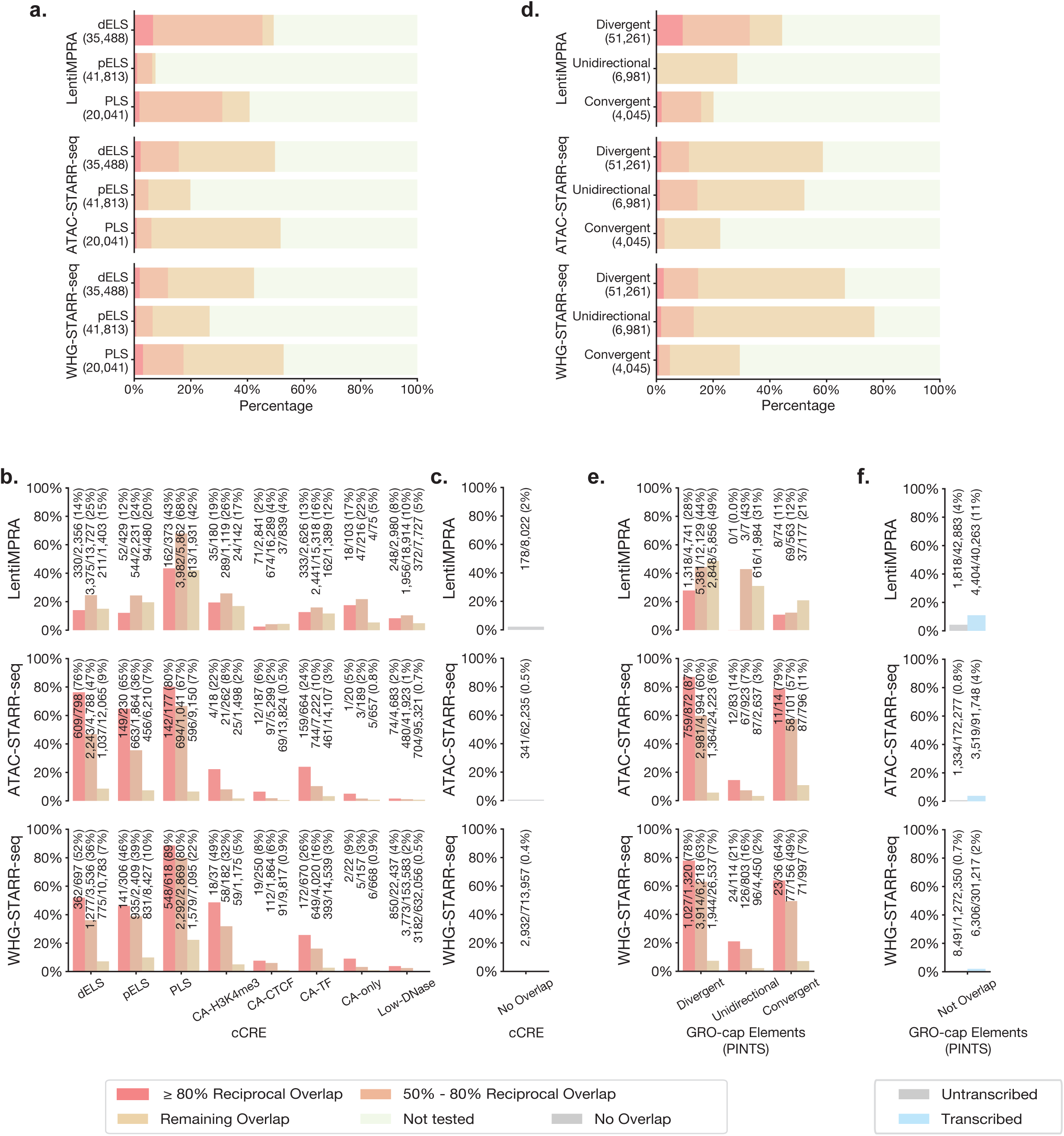
Functional Validation of cCREs and GRO-cap Elements Across MPRA/STARR-seq Assays. **(a)** Stacked bar plot displaying the coverage of dELS, pELS and PLS with different levels of overlap across LentiMPRA, ATAC-STARR-seq and WHG-STARR-seq. Overlap categories include ≥80% reciprocal overlap, 50%-80% reciprocal overlap, and remaining overlap. **(b)** Bar plot illustrating the active rates of all types of cCREs in LentiMPRA, ATAC-STARR-seq and WHG-STARR-seq, stratified by overlap extent. **(c)** Bar plot showing the active rates of all regions not overlapping with any cCREs in LentiMPRA, ATAC-STARR-seq and WHG-STARR-seq. **(d)** Stacked bar plot displaying the coverage of divergent, unidirectional, and convergent elements identified by GRO-cap, categorized by their overlap extent across LentiMPRA, ATAC-STARR-seq and WHG-STARR-seq. **(e)** Bar plot showing the active rates of all types of GRO-cap elements in LentiMPRA, ATAC-STARR-seq and WHG-STARR-seq, stratified by overlap extent. **(f)** Bar plot showing the active rates of regions that do not overlap with GRO-cap elements, stratified by the presence or absence of transcription as measured by GRO-cap signals.

**Extended Data Figure 6.**
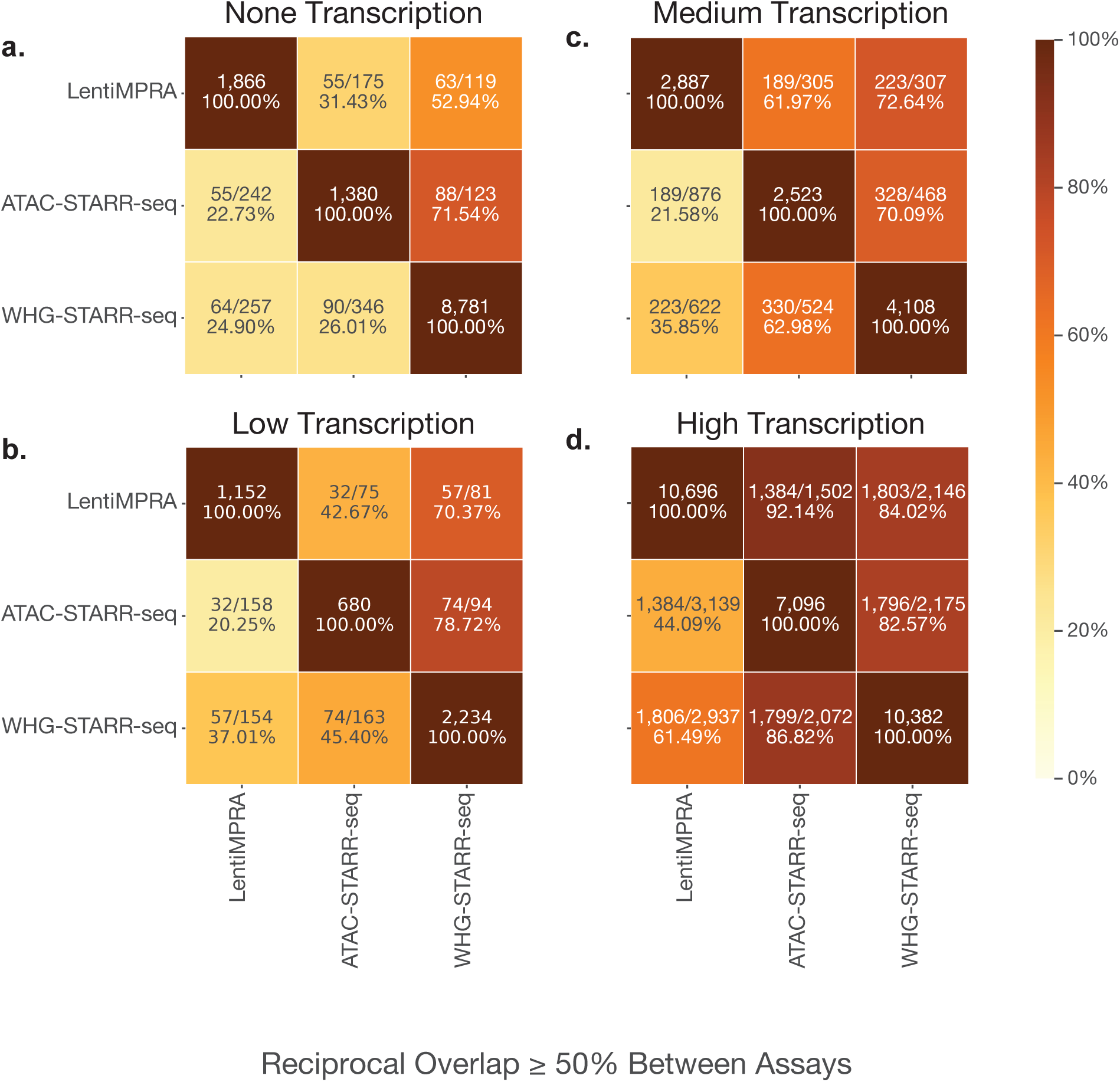
Influence of Transcription Levels on Cross-Assay Comparisons on Enhancer Identification. **(a, b, c,d)** Heatmaps showing the number of commonly active tested regions between assays under the ≥50% reciprocal overlap criterion with none (a), low (b), medium (c) and high (d) transcription levels, as defined by GRO-cap signal. Each cell represents the ratio of the number of enhancer regions in the row dataset that overlap with enhancer regions in the column dataset to the number of enhancer regions in the row dataset overlapping with tested regions in the column dataset. Diagonal cells display the total number of enhancer regions identified in each dataset.

**Supplementary Table 1.**

|  |  | TilingMPRA |  |  | LentiMPRA | ATAC-STARR-seq | WHG-STARR-seq |
| --- | --- | --- | --- | --- | --- | --- | --- |
| Accession |  | ENCSR394HXI | ENCSR917SFD | ENCSR363XER | ENCSR382BVV | ENCSR312UQM | ENCSR661FOW |
| Fragment Size |  | 200-bp | 200-bp | 200-bp | 200-bp | 150-bp - 800-bp | 200-bp - 800-bp |
| Targeted Genomic Regions |  | Tiles in 5-bp sliding window along FEN1, FADS1, FADS2, FADS3 | Tiles in 50-bp sliding window along MYC, GATA1 | Tiles in 100-bp sliding window along LMO2, HBE1, RBM38, HBA2, BCL11A | (1) Protein coding gene promoters centered on TSS; (2) Putative enhancers centered on non-promoter DNase-seq peaks; (3) Tiles not overlapping with DNase peaks around GATA1, MYC, HBE1, LMO2, RBM38, HBA2, and BCL11A | Tn5-fragmented random fragments enriched in accessible regions | Sonicated random fragments genome-wide |
| # DNA Replicates |  | 4 | 5 | 5 | 3 | 6 | 1 |
| # RNA Replicates |  | 4 | 5 | 4 | 3 | 4 | 3 |
| Original Fragments of Interests | # Assayed | 42,714 | 95,990 | 91,110 | 232,542 | 1,136,262,454 (443,451,583) | 826,518,557 |
|  | # Assayed in Forward Orientation | 21,335 | 95,990 | 91,110 | 116,275 | 567,725,494 (221,240,073) | 413,361,239 |
|  | # Assayed in Reverse Orientation | 21,379 | - | - | 116,267 | 568,536,960 (222,211,510) | 413,157,318 |
| # Base Pair Covered |  | 111,020 | 3,912,714 | 9,147,135 | 23,322,354 | 2,816,174,570 (198,059,928) | 2,814,258,311 |
| Control Fragments | # Positive Controls | - | 289 | 290 | 50 | - | - |
|  | # Negative Controls | 150 | 2,871 | 1,874 | 444 | - | - |
| Genomic Bins | # Assayed | - | - | - | - | 557,773,387 (44,507,595) | 549,082,586 |
|  | # Assayed in Forward Orientation | - | - | - | - | 280,006,767 (22,254,224) | 276,158,685 |
|  | # Assayed in Reverse Orientation | - | - | - | - | 277,766,620 (22,253,371) | 282,923,901 |
| % Genome Assayed |  | - | - | - | - | 96.68% (100.00%) | 96.61% |
| # Enhancers (Lab Reported) |  | 13,036 | 17,877 | 14,996 | 87,185 | 53,110 | 38,671 |
| # Enhancers (Merge Overlapping Regions) |  | 121 | 4,419 | 8,379 | 56,840 | 46,906 | 38,671 |

**Supplementary Table 2.**

|  |  | TilingMPRA |  |  | LentiMPRA | ATAC-STARR-seq | WHG-STARR-seq |
| --- | --- | --- | --- | --- | --- | --- | --- |
| Accession |  | ENCSR394HXI | ENCSR917SFD | ENCSR363XER | ENCSR382BVV | ENCSR312UQM | ENCSR661FOW |
| Fragments/Genomic<br>Bins of interests | # Tested | 41,937 | 94,179 | 90,329 | 229,290 | 26,381,426 | 250,455,751 |
|  | # Tested in Both<br>Orientations | 20,611 | - | - | 113,673 | 10,473,380 | 109,842,058 |
| Negative Controls | Source | Originally assayed | Originally assayed | Originally assayed | Originally assayed | Selected from exons | Selected from exons |
|  | # Tested | 148 | 1,828 | 1,862 | 439 | 484,885 | 2,864,061 |
| Z-score Dependent logFC Threshold |  | 2.45 | 1.92 | 2.02 | 0.79 | 1.60 | 1.10 |
| Enhancer Regions | # Either Orientation | 136 | 2,117 | 3,761 | 25,635 | 25,891 | 79,734 |
|  | # Orientation<br>Independent | 57 | - | - | 16,603 | 11,679 | 25,505 |

**Supplementary Table 3.**
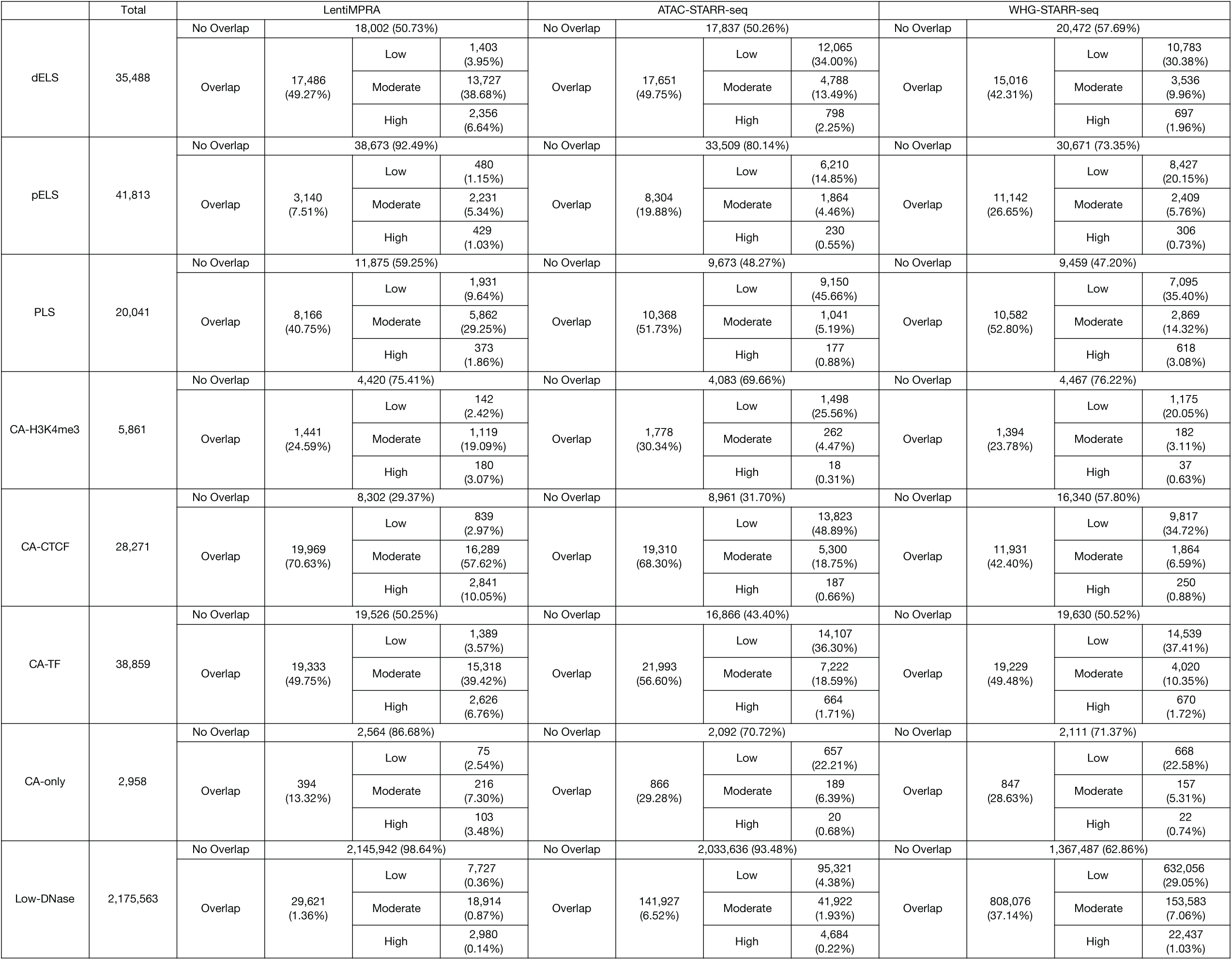

**Supplementary Table 4.**

|  | Total | LentiMPRA |  |  |  | ATAC-STARR-seq |  |  |  | WHG-STARR-seq |  |  |  |
| --- | --- | --- | --- | --- | --- | --- | --- | --- | --- | --- | --- | --- | --- |
| Divergent<br>Elements | 51,261 | No Overlap | 28,535 (55.67%) |  |  | No Overlap | 21,172 (41.30%) |  |  | No Overlap | 17,186 (33.53%) |  |  |
|  |  | Overlap | 22,726<br>(44.33%) | Low | 5,856<br>(11.42%) | Overlap | 30,089<br>(58.70%) | Low | 24,223<br>(47.25%) | Overlap | 34,075<br>(66.47%) | Low | 26,536<br>(51.77%) |
|  |  |  |  | Moderate | 12,129<br>(23.66%) |  |  | Moderate | 4,995<br>(9.74%) |  |  | Moderate | 6,219<br>(12.13%) |
|  |  |  |  | High | 4,741<br>(9.25%) |  |  | High | 871<br>(1.70%) |  |  | High | 1,320<br>(2.58%) |
| Unidirectional<br>Elements | 6,981 | No Overlap | 4,989 (71.47%) |  |  | No Overlap | 3,338 (47.82%) |  |  | No Overlap | 1,614 (23.12%) |  |  |
|  |  | Overlap | 1,992<br>(28.53%) | Low | 1,984<br>(28.42%) | Overlap | 3,643<br>(52.18%) | Low | 2,637<br>(37.77%) | Overlap | 5,367<br>(76.88%) | Low | 4,450<br>(63.74%) |
|  |  |  |  | Moderate | 7<br>(0.10%) |  |  | Moderate | 923<br>(13.22%) |  |  | Moderate | 803<br>(11.50%) |
|  |  |  |  | High | 1<br>(0.01%) |  |  | High | 83<br>(1.19%) |  |  | High | 114<br>(1.63%) |
| Convergent<br>Elements | 4,045 | No Overlap | 3,231 (79.88%) |  |  | No Overlap | 3,134 (77.48%) |  |  | No Overlap | 2,856 (70.61%) |  |  |
|  |  | Overlap | 814<br>(20.12%) | Low | 177<br>(4.38%) | Overlap | 911<br>(22.52%) | Low | 796<br>(19.68%) | Overlap | 1,189<br>(29.39%) | Low | 997<br>(24.65%) |
|  |  |  |  | Moderate | 563<br>(13.92%) |  |  | Moderate | 101<br>(2.50%) |  |  | Moderate | 156<br>(3.86%) |
|  |  |  |  | High | 74<br>(1.83%) |  |  | High | 14<br>(0.35%) |  |  | High | 36<br>(0.89%) |

## Notes

### Competing Interest Statement

The authors have declared no competing interest.

